# Decoding social integration in schooling fish using real–virtual interactions

**DOI:** 10.64898/2026.03.12.711426

**Authors:** Zhen Kang, Ramón Escobedo, Maud Combe, Stéphane Sanchez, Clément Sire, Guy Theraulaz

**Affiliations:** School of Systems Science, Beijing Normal University, 100875 Beijing, China; Centre de Recherches sur la Cognition Animale, Centre de Biologie Intégrative, CNRS, Université de Toulouse III – Paul Sabatier, 31062 Toulouse, France; Laboratoire de Physique Théorique, CNRS, Université de Toulouse III – Paul Sabatier, 31062 Toulouse, France; Institut de Recherche en Informatique de Toulouse (IRIT), Université Toulouse Capitole, Toulouse, France; Departamento de Matemáticas, Universidad Carlos III de Madrid, Leganés, Madrid, Spain

**Keywords:** Collective motion, Virtual reality, Fish school model, 3D agent-based modeling, Social interactions, Closed loop

## Abstract

Collective motion in animal groups emerges from local interactions, yet how individuals select and integrate social information remains poorly understood. In schooling fish, inferring which neighbors drive an individual’s behavior at any given moment is extremely challenging because the analysis of trajectories alone does not unambiguously reveal the underlying causal interactions that generate the observed motion. Here, we used a closed-loop virtual-reality system in which a single real rummy-nose tetra *Hemigrammus rhodostomus* interacted in real time with four model-driven virtual conspecifics whose interaction rules were precisely controlled. By systematically varying the number of influential neighbors of the virtual fish and combining experiments with simulations, we quantified how social information filtering shapes individual and collective dynamics. Collective coordination increased with stronger social coupling, but analyses of kinematics, spatial organization, correlations, and turning dynamics consistently showed that the real fish behaved as if it were responding primarily to the single most influential neighbor, whose identity could change over time. These results demonstrate that selective interaction with the most influential neighbor is sufficient to sustain coordinated group motion and highlight the power of bio-hybrid closed-loop experiments to reveal causal mechanisms of collective behavior.

## I. INTRODUCTION

Collective motion in animal groups is one of the most striking manifestations of self-organization in living systems [1, 2]. Schools of fish, flocks of birds, herds of ungulates, and swarms of insects display highly coordinated movements that emerge from local interactions among individuals without centralized control [3]. Over the past several decades, theoretical and empirical work has demonstrated that such patterns can arise from relatively simple interaction rules based on attraction, repulsion, and alignment with neighbors [2, 4–9]. These findings have established collective motion as a paradigmatic example of emergent behavior in complex systems and have stimulated a broad interdisciplinary effort spanning behavioral biology, physics, and computational modeling to understand how individual-level decisions scale up to group-level organization and how individuals perceive, process, and respond to social information [2, 10–12].

Despite this progress, a fundamental question remains unresolved: how do individuals in a moving group combine and integrate the information coming from their neighbors in order to coordinate their own movements? This question is central because the way social information is filtered and integrated at the individual level ultimately determines the macroscopic properties that emerge at the collective level, including the stability of coordination, the speed of information propagation, and the responsiveness of the group to environmental perturbations.

Theoretical studies have shown that collective dynamics can depend sensitively on the number of influential neighbors and on the rules governing their selection, with excessive information sometimes degrading rather than improving coordination [13–17]. At the same time, converging empirical evidence indicates that animals do not process all available social cues indiscriminately, but instead rely on selective and context-dependent interactions shaped by perceptual, cognitive, and sensory constraints. A growing body of work suggests that collective coordination often relies on the selective rather than exhaustive integration of social information, with individuals attending preferentially to a limited subset of neighbors, sometimes even to a single influential individual, depending on spatial configuration, motion cues, or behavioral salience [18–21]. Such parsimonious interaction strategies reduce the cognitive and perceptual load while preserving robust collective coordination, indicating that filtering of social information is not merely a constraint but an adaptive feature of distributed decision making in animal groups.

A major challenge in addressing these issues lies in the difficulty of identifying the actual interaction rules used by real animals. Advances in automated tracking and quantitative analysis of trajectories have made it possible to infer candidate behavioral rules by examining how individuals adjust their motion in response to the positions and movements of neighbors [5, 6, 8, 9]. However, similar collective patterns can arise from different microscopic rules, and statistical agreement between simulations and experiments does not guarantee that the inferred behavioral processes are correct [8, 22, 23]. Establishing causal links between hypothesized interaction rules and observed behavior therefore requires experimental approaches that allow the systematic manipulation of social stimuli while preserving natural sensorimotor feedback.

In recent years, closed-loop virtual reality systems have emerged as a powerful methodology to address this limitation. By embedding animals in interactive environments where visual stimuli are dynamically updated according to their own movements, virtual reality makes it possible to control specific aspects of the social environment with high precision while maintaining ecological realism [24–29]. In such closed-loop systems, animals interact with virtual conspecifics that respond in real time, enabling direct tests of behavioral models and hypotheses about social integration and coordination. Previous work has shown that fish respond to virtual partners in ways that closely resemble interactions with real conspecifics, adjusting their speed, position, and orientation according to the behavior of virtual agents [29, 30]. These findings demonstrate that virtual stimuli can elicit biologically meaningful social responses and provide a foundation for using virtual reality to investigate the mechanisms underlying collective behavior.

In this study, we address directly the question of how many neighbors a fish effectively uses to control its swimming behavior while moving in a group and how filtering social information shapes collective dynamics. Experiments were conducted with rummy-nose tetra *Hemigrammus rhodostomus*, a species exhibiting strong schooling behavior and widely used as a model system for quantitative studies of collective motion because of its robust polarization, cohesive group structure, and reproducible social interactions [9, 31–33]. We investigated bio-hybrid groups composed of one real fish interacting in real time with four virtual conspecifics whose behavior was governed by a 3D data-driven model of social coordination. This experimental paradigm makes it possible to manipulate precisely the number of influential neighbors affecting the motion of the virtual fish while keeping all other conditions constant, thereby providing an experimental handle on a question that is otherwise difficult to test in natural groups.

By combining closed-loop experiments with predictive modeling, our study aims to bridge the gap between statistical inference of interaction rules and their causal validation. More broadly, understanding how individuals filter and integrate social information is essential for explaining how collective behavior emerges in biological systems and for designing artificial collectives capable of robust decentralized coordination [11, 34]. The results reported here contribute to this objective by providing quantitative evidence on the mechanisms of social integration in schooling fish and by illustrating how controlled bio-hybrid experiments can reveal the causal structure of collective behavior.

## II. RESULTS

We examined two related issues. First, we asked how the collective behavior of the group depends on the number of influential neighbors that the virtual fish take into account. To answer this question, we combined experiments and model simulations, exploiting the same threedimensional behavioral model (see Materials and Methods) to command the virtual fish in the experiments and to command the real fish and the virtual fish in the simulations. In the model, the pairwise interactions of a focal agent with each other agent are first computed and ranked according to their magnitude. Then, for a given value of *k* for the focal agent, the actual acceleration of the focal agent is computed by only summing the contributions of the *k* neighbors corresponding to the *k* largest pairwise interactions. The *k* neighbors selected in this manner are dubbed the *k* (most) influential neighbors of the focal agent at a given time [20]. Note that their actual identities change with time as the group continuously reorganizes. In the experiments, we systematically varied the number of influential neighbors of the virtual fish (*k*_V_ = 1, 2, and 4), while the real fish interacted freely with them. In the simulations, we varied not only the number of influential neighbors of the virtual fish (*k*_V_) but also the number of influential neighbors of the simulated real fish (*k*_R_ = 1, 2, and 4).

Second, we examined how these interaction rules affect the behavior of real and virtual individuals separately. By analyzing the responses of the real fish and the virtual fish independently in both experiments and simulations, we were able to determine which assumptions regarding the effective number of neighbors used by the real fish best reproduce the observed dynamics.

This joint comparison of experiments and simulations allowed us to determine which hypothesis regarding *k*_R_ best reproduces the observed collective dynamics and to infer the effective number of neighbors used by the real fish to coordinate its motion. The experimental setup and procedures are described in detail in the Materials and Methods section.

Figure 1A illustrates the bio-hybrid closed-loop interaction system used in this study, in which a real fish swims in a hemispherical arena while interacting in real time with four virtual conspecifics whose behavior is continuously updated by our behavioral model in response to the movements of the real fish (see also Supplementary Movies 1, 2, 3 and 4). Figure 1B summarizes the key kinematic and spatial variables used to describe and quantify individual motion and the effects of social interactions on swimming behavior. Precise definitions and mathematical formulations of these variables, together with those used to fully characterize individual and collective behavior, are provided in the Materials and Methods section.

**FIG. 1.**
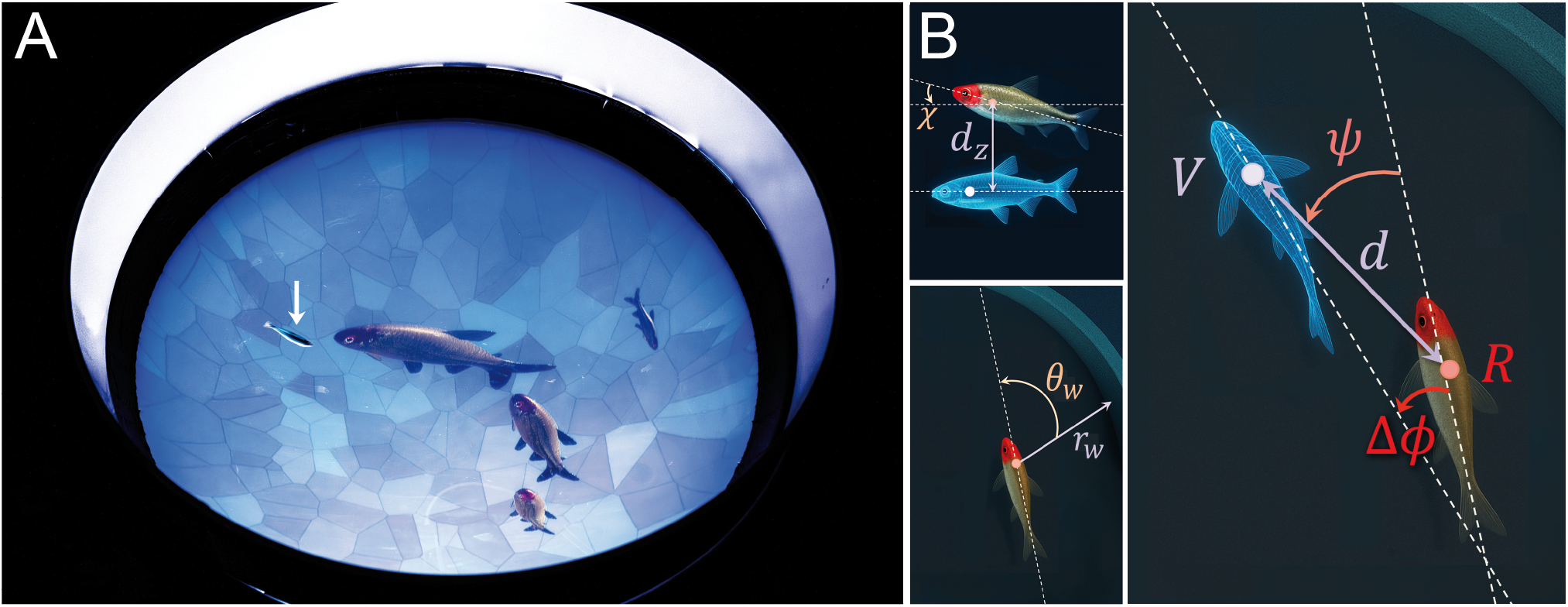
Closed-loop 3D virtual-reality system and interaction geometry. (A) Hemispherical bowl in which a real fish (white arrow) swims while interacting with four virtual conspecifics displayed as anamorphic projections. The behavior of the virtual fish is controlled by a mathematical model that updates their positions and orientations in real time in response to the movements of the real fish. (B) Schematic representation of the variables used for behavioral analysis. These include the elevation angle *χ* of the real fish (red) relative to the horizontal plane, the vertical separation between fish *d*_z_, the distance from the wall *r*_w_ and the angle of incidence to the wall *θ*_w_ in the horizontal plane, the horizontal distance between fish *d*, the viewing angle *ψ* at which the real fish (red, R) perceives a virtual conspecific (blue, V), and the relative orientation Δ*ϕ* = *ϕ*_V_ *−ϕ*_R_, where *ϕ* denotes the azimuthal heading angle of each fish, defined from the projection of the velocity vector onto the horizontal plane (direction indicated by dashed lines).

The behavioral model used to control the four virtual fish in the experiments was identical to the one employed in the numerical simulations, as noted above. For each social interaction strategy (*k* = 1, 2, and 4, identical for all agents), model parameters were calibrated to satisfactorily reproduce the experimental results obtained in groups of five fish swimming in a circular arena (see Fig. S1). This calibration ensured quantitative agreement between simulation outcomes and empirical distributions of individual and pairwise behavioral variables. The full calibration procedure is described in the Materials and Methods section.

### A. Social-information filtering shapes collective dynamics in bio-hybrid fish groups

We first investigated how filtering social information by limiting the number of influential neighbors shapes the collective dynamics of bio-hybrid groups composed of one real fish and four virtual conspecifics. Using experiments and model simulations, we compared conditions in which virtual fish interacted with only their most influential neighbor (*k*_V_ = 1), their two most influential neighbors (*k*_V_ = 2), or all four neighbors (*k*_V_ = 4).

Representative trajectories reveal a clear effect of the number of influential neighbors on group coordination (Fig. 2), with increasing *k*_V_ leading to progressively stronger coordination, as indicated by a systematic rise in mean polarization. Notably, only the configuration in which both the virtual fish and the simulated real fish interacted exclusively with their single most influential neighbor qualitatively reproduced the experimental dynamics observed for *k*_V_ = 1. Simulations with larger values of *k*_R_ produced more coherent and stable trajectories, indicating that even a single individual integrating information from multiple neighbors can enhance global coordination (Fig. S2).

**FIG. 2.**
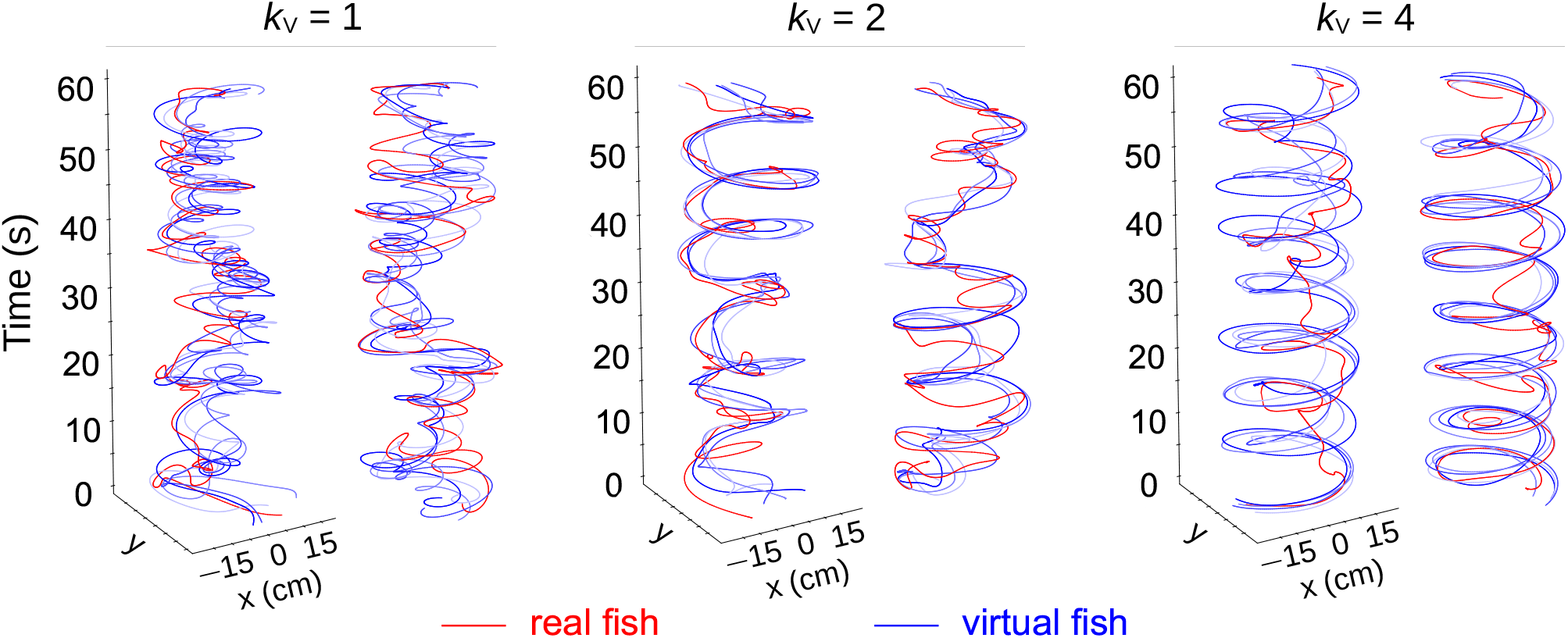
Collective movement patterns under different social interaction strategies of the virtual fish. Samples of trajectories of the real fish (red) and the four virtual conspecifics (blue shades) recorded during six experimental sessions for three different values of the number of influential neighbors considered by the virtual fish: *k*_V_ = 1, *k*_V_ = 2, and *k*_V_ = 4. Two representative one-minute trajectory segments are shown for each condition.

Both experiments and simulations revealed transient departures of the real fish from the virtual group, followed by rapid reintegration (see Supplementary Movies 4, 5, 6 and 7). These events were particularly frequent when *k*_V_ = 2 or 4 and when *k*_R_ was small (Fig. 2, Fig. S2). As *k*_R_ increased, such departures became rare and were nearly absent for *k*_R_ = 4, indicating that integrating information from multiple neighbors stabilizes group cohesion.

At the collective level, the probability density functions of polarization and milling captured the emergence of coordinated states (Fig. 3; Supplementary Table 3). Experimental results were closely reproduced by the model when *k*_V_ = 1, independently of the value of *k*_R_. Fig. S3A shows that the cumulative distribution function (CDF) of polarization when *k*_V_ = 2 is consistently below that of *k*_V_ = 1, and further below for *k*_V_ = 4 compared to *k*_V_ = 2, indicating higher polarization for larger values of *k*_V_. A similar trend is observed in simulations for all *k*_R_ = 1, 2, 4 (Fig. S3B–D).

**FIG. 3.**
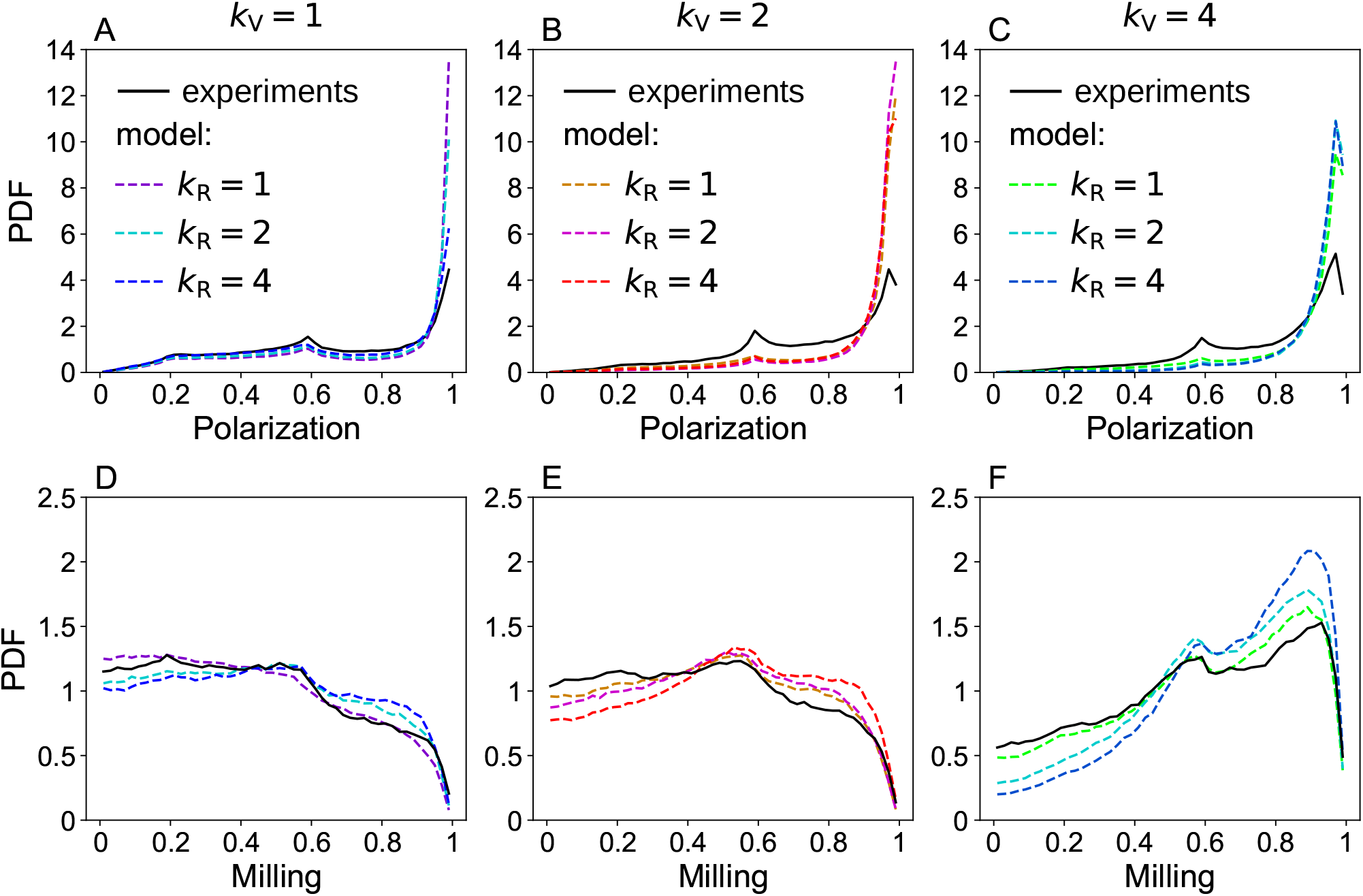
Collective order in bio-hybrid groups composed of one real fish and four virtual conspecifics under different social interaction strategies. Probability density functions (PDFs) of polarization (A–C) and milling (D–F) for the three social interaction regimes of the virtual fish, *k*_V_ = 1, 2, and 4. Black solid lines correspond to experimental data. Colored dashed lines represent model simulations for each value of *k*_V_ and for the three social interaction strategies of the real fish, *k*_R_ = 1, 2, and 4.

Occasional configurations in which one fish moved opposite to the others produced a secondary peak near *P≈* 0.6, consistent with theoretical expectations for a group of five individuals. Such configurations were rare in simulations when virtual fish interacted with multiple neighbors, indicating that increasing *k*_V_ stabilizes aligned states.

When *k*_V_ = 4, polarization and milling increased jointly, reflecting the fact that individuals moved as a cohesive group while remaining close to the circular boundary of the arena. For *k*_V_ = 2 and 4, both polarization and milling also increased with *k*_R_ in the simulations. However, the values measured in the experiments remained substantially lower than those predicted by the model for bio-hybrid groups in which all individuals would interact with the same number of neighbors. This discrepancy suggests that under these conditions, real fish effectively interact with fewer neighbors than assumed in the corresponding simulations.

Individual kinematics remained remarkably stable across conditions. Groups swam at a preferred speed of v *≈* 10–11 cm s^*−*1^ (Fig. 4A, D, G) and remained close to the bowl wall, with the mean wall distance decreasing as *k*_V_ increased (Fig. 4B, E, H). Accordingly, the PDFs of the heading angle relative to the wall *θ*_w_ exhibit peaks at *±*90^*°*^ whose magnitude increases with *k*_V_, indicating increasingly tangential motion along the boundary (Fig. 4C, F, I, where the peaks are twice as high for *k*_V_ = 4 as for *k*_V_ = 1). Model simulations quantitatively reproduced these patterns, and varying *k*_R_ had little influence on speed, wall distance, or incidence angle, as confirmed by the Hellinger distance quantifier (see Supplementary Table 2 and Materials and Methods). Fish also exhibited a preferred swimming depth around z*≈* 4.5 cm, with occasional excursions toward the bottom of the tank (Fig. S4A–C).

**FIG. 4.**
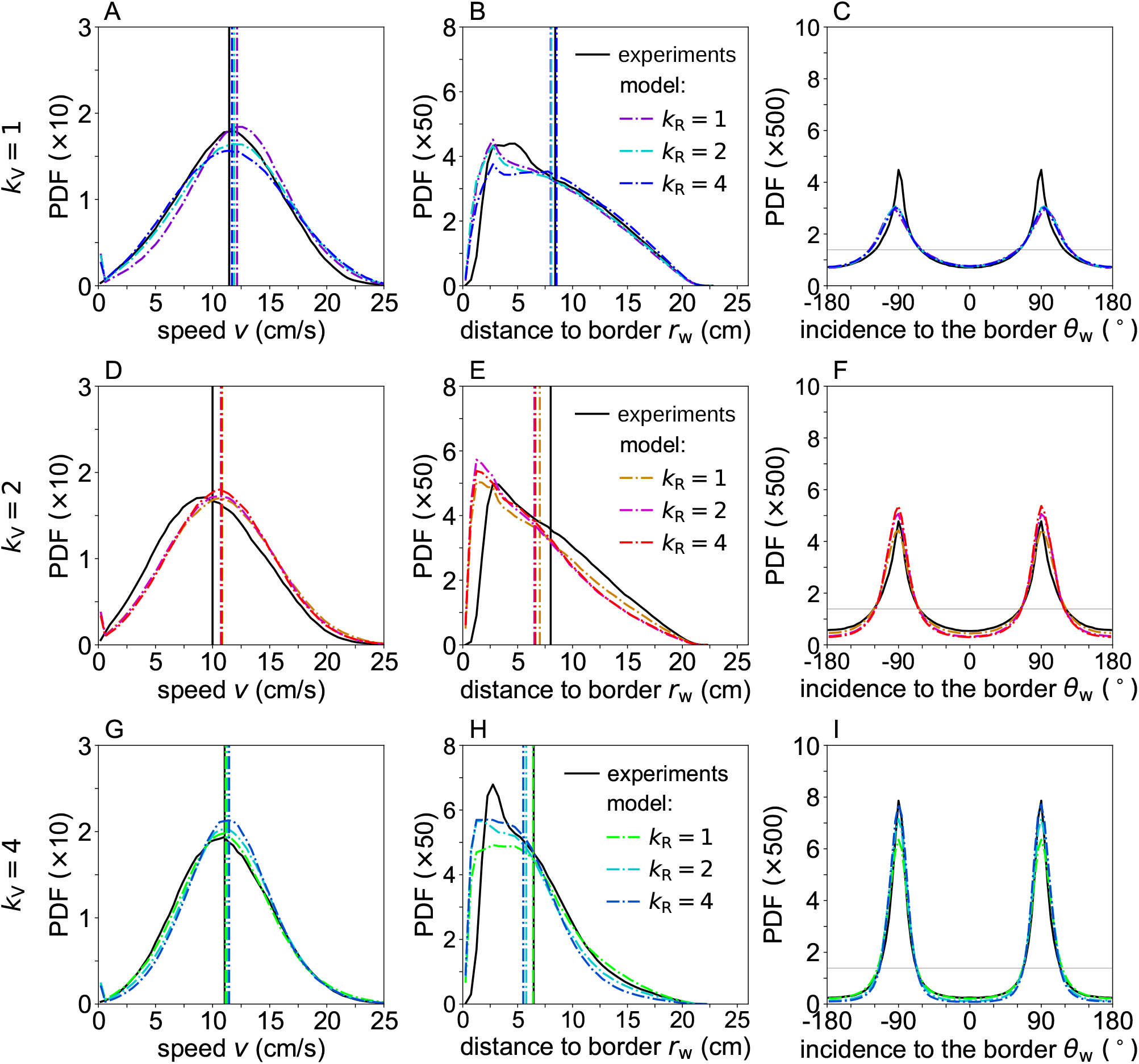
Individual kinematics and wall-interaction statistics in bio-hybrid groups under different social interaction strategies. Probability density functions (PDFs) of (A, D, G) individual swimming speed v, (B, E, H) distance to the wall *r*_w_, and (C, F, I) angle of incidence to the wall *θ*_w_, for *k*_V_ = 1 (first row), *k*_V_ = 2 (second row), and *k*_V_ = 4 (third row). Black solid lines correspond to experimental data. Colored dashed lines represent model simulations for each value of *k*_V_ and for the three social interaction strategies of the real fish, *k*_R_ = 1, 2, and 4. Vertical lines indicate the mean value of the corresponding PDF in matching colors. The horizontal grey lines in panels (C, F, I) denote the uniform distribution.

Spatial organization within the group also depended on the number of influential neighbors. Individuals remained in close proximity, with typical horizontal separations around *d ≈*9 cm and small vertical separations (Fig. 5A, D, G; and Fig. S4D–F). Relative positions were weakly structured when *k*_V_ = 1, but became increasingly stable as *k*_V_ increased. In these conditions (Fig. 5B, E, H), fish most often viewed their nearest neighbor directly ahead or behind, and heading differences were strongly peaked around zero, revealing robust alignment.

**FIG. 5.**
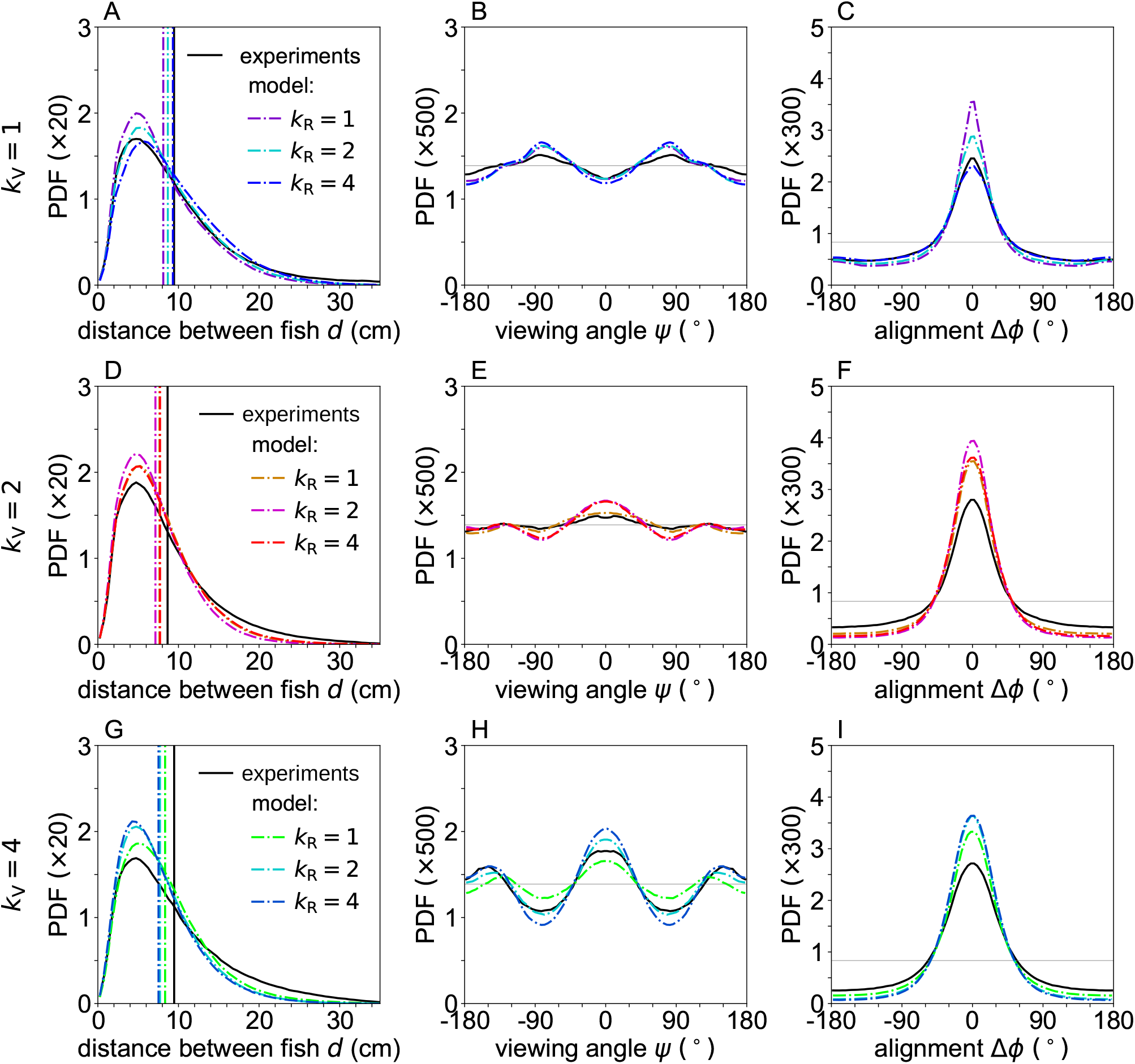
Pairwise spatial and orientational interaction statistics in bio-hybrid groups under different social interaction strategies. Probability density functions (PDFs) of (A, D, G) distance between fish *d*, (B, E, H) viewing angle *ψ*, and (C, F, I) heading-angle difference Δ*ϕ*, for *k*_V_ = 1 (first row), *k*_V_ = 2 (second row), and *k*_V_ = 4 (third row). Black solid lines correspond to experimental data. Colored dashed lines represent model simulations for each value of *k*_V_ and for the three social interaction strategies of the real fish, *k*_R_ = 1, 2, and 4. Vertical lines in panels (**A, D, G**) indicate the mean value of the corresponding PDF in matching colors. Horizontal grey lines in the angular PDFs denote the uniform distribution.

Model simulations qualitatively reproduced the influence of *k*_V_ on horizontal inter-individual distance *d*, viewing angle *ψ*, and heading alignment Δ*ϕ*. Increasing *k*_R_ further promoted a stable front–back aligned and coherent swimming state, clearly visible in Fig. 5H, together with increased alignment between neighbors.

Together, these results show that filtering social information by limiting the number of interacting neighbors strongly shapes collective dynamics. Larger values of *k*_V_ produce clear and systematic effects, enhancing alignment and coordination, whereas increasing *k*_R_ mainly stabilizes cohesion and coherent motion. For each value of *k*_V_, all three values of *k*_R_ yielded statistically indistinguishable agreement to the experimental data, as quantified by the Hellinger distances (Supplementary Table 2). Thus, collective-level metrics alone do not allow us to discriminate between alternative hypotheses regarding the number of neighbors effectively used by the real fish. Resolving this ambiguity requires analyzing the behavior of real and virtual fish separately. We turn to this analysis next.

### B. Real-fish responses reveal effective single-neighbor interactions in bio-hybrid groups

We next examined how the behavior of the real fish depends on the level of coordination among virtual conspecifics. Figure 6 shows the average individual responses of the real and virtual fish in the experiments. Across all conditions, the real fish swam systematically faster than the virtual fish (Fig. 6A, D, G; Supplementary Table 3). As *k*_V_ increased, both real and virtual fish moved closer to the wall and aligned more strongly with it (Fig. 6B, E, H, and C, F, I; Supplementary Table 4), reflecting the increasing coordination of group motion. These effects are also clearly visible in the trajectories shown in Fig. 2, where higher values of *k*_V_ lead to smoother, more coherent paths and a tighter spatial organization of the group.

**FIG. 6.**
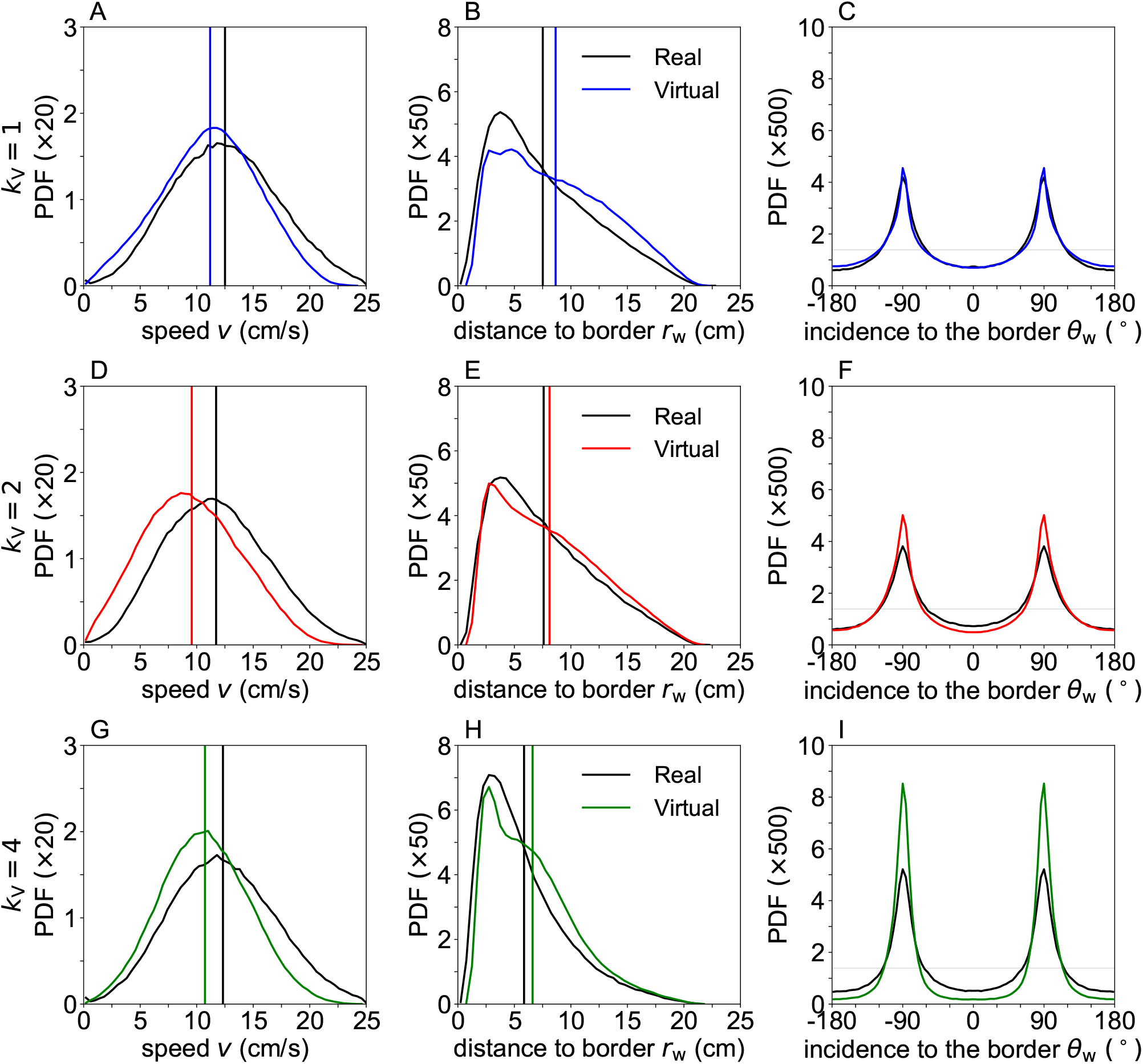
Comparison of the kinematics and wall-interaction statistics of real fish (black lines) and averaged virtual conspecifics (colored lines) under different social interaction strategies of the virtual fish in experiments. Probability density functions (PDFs) of (A, D, G) individual swimming speed v, (B, E, H) distance to the wall *r*_w_, and (C, F, I) angle of incidence to the wall *θ*_w_, for *k*_V_ = 1 (first row), *k*_V_ = 2 (second row), and *k*_V_ = 4 (third row). Vertical lines indicate the mean value of the corresponding PDF in matching colors. Horizontal grey lines in panels (**C, F, I**) denote the uniform distribution.

Despite this global trend, the responses of the real fish differed from those of the virtual fish in a systematic way. When *k*_V_ = 1, the alignment of the real fish with the wall was similar to that of the virtual fish (Fig. 6C). At larger values of *k*_V_, however, alignment increased only weakly in the real fish, whereas it continued to increase for the virtual fish (Fig. 6F, I). This difference suggests that the real fish integrates social information over a smaller number of neighbors. Because the identity of the most influential neighbor may change from one orientation update to the next, the real fish exhibits larger directional fluctuations, visible as a broader distribution of *θ*_w_ around *±*90^*°*^, particularly for *k*_V_ = 4.

Changes in inter-individual distances further support this interpretation. The mean distance between the real fish and the virtual group increased with *k*_V_ (Fig. 7A, D, G; Supplementary Table 4), from approximately 10.0 cm at *k*_V_ = 1 to 11.6 cm at *k*_V_ = 4, while the group of virtual fish became slightly more compact. However, the distance between the real fish and its nearest virtual neighbor remained approximately constant and is similar to the distance measured in pairs of real fish swimming together. This value is consistent with previous results showing that the attractive interaction between fish decreases sharply at short range, so that individuals rarely approach each other below a distance of about 5 cm [30].

**FIG. 7.**
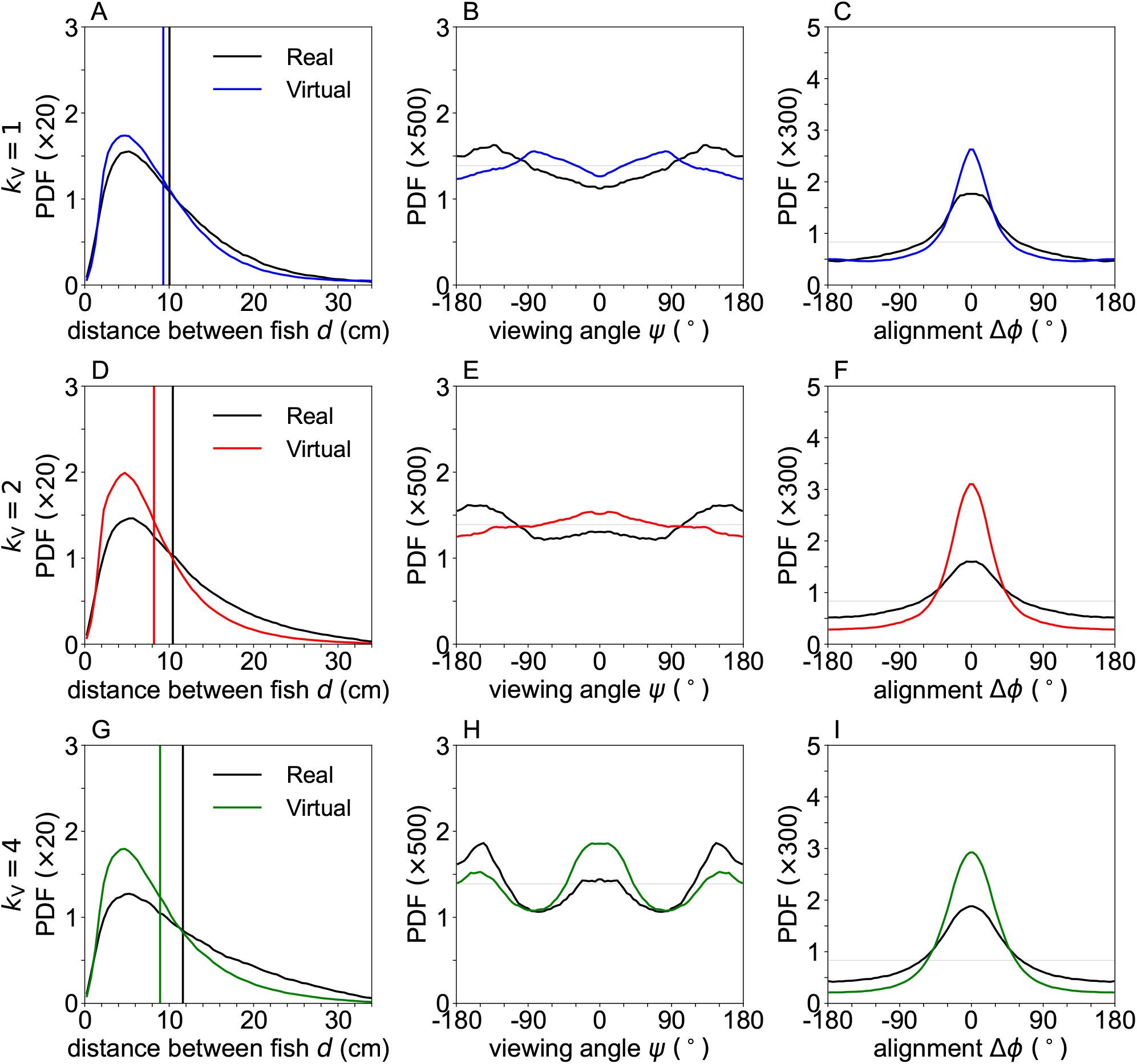
Comparison of the pairwise interaction geometry between real fish (black lines) and averaged virtual conspecifics (colored lines) under different social interaction strategies of the virtual fish in experiments. Probability density functions (PDFs) of (A, D, G) inter-individual distance *d*, (B, E, H) viewing angle *ψ*, and (C, F, I) headingangle difference Δ*ϕ*, for *k*_V_ = 1 (first row), *k*_V_ = 2 (second row), and *k*_V_ = 4 (third row). Vertical lines in panels (A, D, G) indicate the mean value of the corresponding PDF in matching colors. Horizontal grey lines in the angular PDFs denote the uniform distribution.

Vertical organization showed a similar pattern. When *k*_V_ = 1, the real and virtual fish swam at similar depths and maintained comparable vertical separations (Fig. S5; Supplementary Tables 4 and 5). As *k*_V_ increased, real fish tended to swim slightly deeper and to remain marginally farther from the virtual group, consistent with weaker coupling to the collective motion.

The spatial organization of interactions was further revealed by the distributions of viewing angles (Fig. 7). For *k*_V_ = 1, the distribution of the viewing angle of the real fish to virtual conspecifics was relatively flat, with weak peaks around *±*135^*°*^, indicating limited spatial structure and a tendency for the real fish to be located in front of the virtual group along the direction of motion. When positioned in front, the real fish may transiently act as an incidental leader due to social interactions with the virtual fish, although this spatial configuration alone does not imply sustained leadership. As *k*_V_ increased, an additional peak appeared near 0^*°*^, showing that the real fish also spent substantial time following virtual conspecifics. Virtual fish exhibited a similar but more pronounced spatial organization, spending more time following neighbors, as reflected by stronger peaks near 0^*°*^.

Consistent with these patterns, increasing coordination among virtual fish reduced the frequency with which the real fish temporarily left the group and performed spontaneous U-turns. The time intervals between successive U-turns increased with *k*_V_ (Fig. S6), indicating that stronger collective order stabilizes the motion of the real fish.

Cross-correlation analyses provided further insight into interaction dynamics. Correlations between the velocity vector of the real fish and those of virtual fish showed the closest agreement between experiments and simulations when *k*_R_ = 1 (Fig. S7A, C, E). In experiments, the correlation peaked at time lags of 0.54 s for *k*_R_ = 1, 0.8 s for *k*_R_ = 2, and 0.34 s for *k*_R_ = 4. By contrast, correlations among virtual fish were strong and symmetric for all values of *k*_V_, reflecting their tightly coordinated motion (Fig. S7B, D, F).

Turning dynamics provided one of the clearest discriminating signatures between interaction hypotheses. The probability density functions of the signed angular speed *dϕ*^+^/*dt* show that real fish frequently perform large-amplitude turns, producing a pronounced high-value tail in the experimental distributions across all values of *k*_V_ (Fig. 8). In particular, angle changes larger than 60^*°*^, and even exceeding 90^*°*^, occur regularly in the experiments but are reproduced in simulations only when the real fish is assumed to interact with a single influential neighbor (*k*_R_ = 1). A second distinctive feature concerns the direction of turning. Turns directed toward the wall, corresponding to negative values of *dϕ*^+^/*dt*, are observed experimentally with substantial frequency but are again reproduced only in simulations with *k*_R_ = 1. When the simulated real fish integrates information from two or four neighbors, both large turns and turns toward the wall become rare or disappear altogether, especially for *k*_V_ = 2 and 4. These results indicate that increasing the number of neighbors taken into account in the decision process effectively reduces maneuverability and smooths directional changes. Most importantly, the turning statistics provide a clear behavioral indicator that the dynamics of the real fish are best explained when it responds to a single most influential neighbor, independently of the interaction strategy of the virtual fish.

**FIG. 8.**
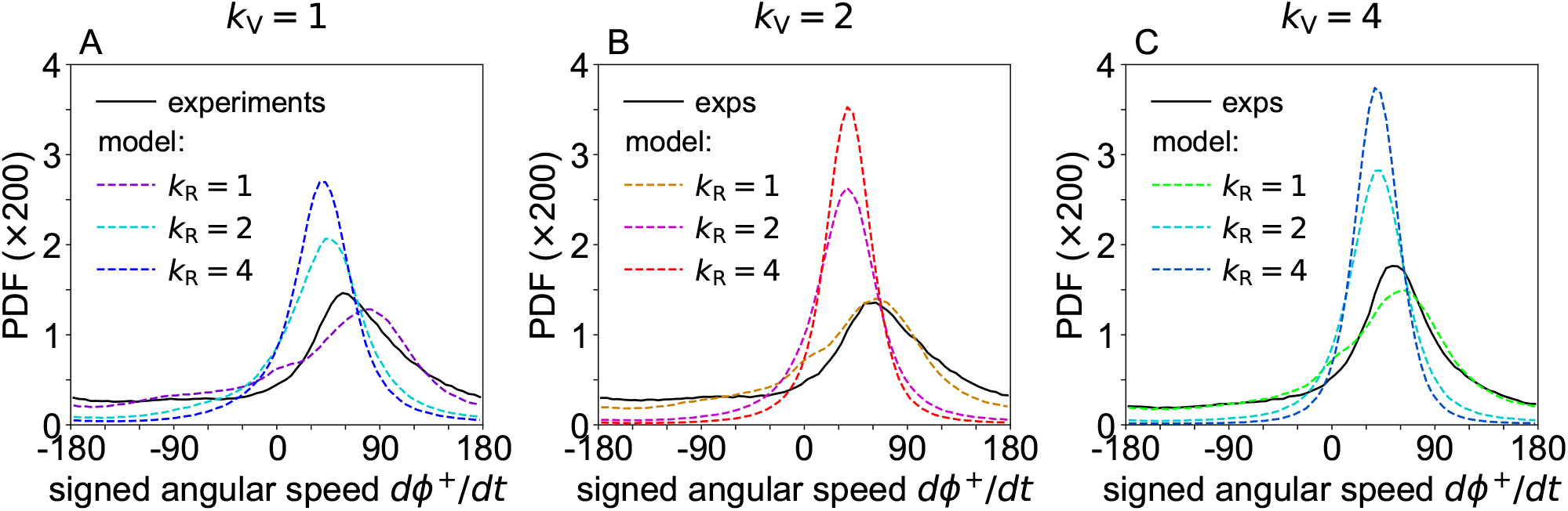
Turning kinematics of real fish under different social interaction strategies. Kinematics of turning maneuvers of the real fish, quantified by the probability density functions (PDFs) of the signed angular velocity *dϕ*^+^*/dt* (sign determined by the wall-incidence angle *θ*_w_), for (A) *k*_V_ = 1, (B) *k*_V_ = 2, and (C) *k*_V_ = 4. Black solid lines correspond to experimental data. Colored dashed lines represent model simulations for each value of *k*_V_ and for the three social interaction strategies of the real fish, *k*_R_ = 1, 2, and 4.

Finally, analyses of the distances and spatial positions of the most influential neighbors (Figs. S8–S10) showed that the most influential neighbors of the real fish are not always the nearest ones and can occupy a wide range of spatial positions. These results also confirm that the virtual reality system faithfully reproduces realistic spatial configurations of virtual conspecifics within the visual field of the real fish.

Taken together, all behavioral, spatial, and dynamical indicators consistently converge toward the same conclusion: real fish effectively coordinate their movements by interacting with only their most influential neighbor.

## III. DISCUSSION

How animals combine social information to coordinate their movements remains a central question in the study of collective behavior [8, 16, 35]. This integration process determines how local interactions scale up to group-level patterns such as alignment, cohesion, and coordinated turning. Yet, identifying how many neighbors an individual effectively uses is difficult in natural settings, where interactions are continuous and strongly coupled. Moreover, most empirical approaches infer interaction rules indirectly from statistical similarities between simulations and experiments, a strategy that cannot fully establish the causal mechanisms of social coordination [36]. More broadly, determining the causal structure of interaction networks in mobile animal groups remains one of the major methodological challenges in collective behavior research, because observational data alone rarely allow discrimination between alternative hypotheses about social influence or sensory integration [19, 27, 37].

Here we addressed this problem using a bio-hybrid closed-loop approach in which a single real fish interacted in real time with four virtual conspecifics. The behavior of the virtual fish was governed by a validated three-dimensional swimming model whose interaction parameters were calibrated to reproduce previous experiments with groups of five real fish. Because the behavior of the virtual fish was fully controlled, we could manipulate the number of influential neighbors shaping their motion while keeping all other conditions constant. This approach provided an experimental handle on a question that is otherwise difficult to test in natural groups. Closed-loop virtual reality experiments provide a particularly powerful framework in this respect because they allow reciprocal interactions and therefore enable direct behavioral validation of interaction rules rather than relying solely on statistical similarity between model outputs and empirical distributions [24, 27, 30].

We defined *k* as the number of most influential neighbors considered when a fish updates its heading. In the experiments, we systematically varied the number of influential neighbors of the virtual fish (*k*_V_ = 1, 2, or 4). In the model simulations, we reproduced these experimental conditions by using the same values of *k*_V_ and, for each of these three conditions, we further assumed that the real fish integrated information from one, two, or four neighbors (*k*_R_ = 1, 2, or 4). This framework builds on previous work showing that realistic schooling dynamics can be reproduced by combining experimentally reconstructed attraction and alignment functions with stochastic burst-and-coast locomotion, providing a quantitative bridge between behavioral analyses and predictive models [9, 20, 38]. This factorial design allowed us to compare experiments and simulations directly and to determine which hypothesis about *k*_R_ best accounts for the observed behavior.

Our first result is that filtering social information by limiting the number of influential neighbors strongly shapes collective dynamics. Larger values of *k*_V_ systematically increased polarization and coordination in the bio-hybrid groups. Groups became more cohesive, fish trajectories became smoother, and departures of the real fish from the virtual subgroup became less frequent. These effects were robust across experiments and simulations, indicating that the number of influential neighbors is a key control parameter of collective motion. This sensitivity to interaction filtering aligns with theoretical and computational studies showing that collective order depends critically on how individuals weight and select social information, and that selective interaction rules can regulate transitions between disordered and ordered states [13, 15, 17, 39].

A striking pattern emerged when *k*_V_ = 4. In this condition, polarization and milling increased simultaneously. Fish moved as a cohesive group while remaining close to the circular boundary of the arena. In such conditions, tangential motion along the wall naturally promotes rotational dynamics. This result shows that increasing social coupling does not only stabilize alignment; it can also stabilize coherent circulation depending on environmental constraints. The coexistence of multiple collective states under strong coupling echoes predictions from physical and biological models in which interaction topology, sensory constraints, and boundary interactions jointly shape phase transitions and multistability in collective motion [2, 37, 39, 40].

A second important finding concerns the role of *k*_R_. In simulations, increasing the number of neighbors integrated by the real fish stabilized trajectories and reduced transient departures. This result is intuitive, since averaging information from multiple neighbors reduces directional noise and produces smoother motion. However, experimental levels of polarization and milling remained consistently lower than those predicted by simulations in which all individuals used large values of *k*. This discrepancy suggests that real fish effectively use fewer neighbors than assumed in these simulations. Experimental and modeling studies of visual attention and decision-making indicate that fish often select one target at a time, effectively truncating information from other neighbors, a strategy that can naturally produce the type of variability observed in our experiments [14, 21, 41].

A third key point is that collective-level observables alone were not sufficient to identify the effective number of neighbors used by the real fish. For each value of *k*_V_, simulations with different values of *k*_R_ produced statistically indistinguishable results for most global metrics. Only the variables reflecting spatial organization, such as viewing-angle distributions and directional variability, provided a clear discrimination. This result highlights the limits of relying exclusively on global order parameters when inferring interaction rules and supports the growing recognition that the topology and dynamics of interaction networks must be reconstructed explicitly to understand behavioral contagion and influence patterns in mobile groups [5, 19, 28].

Individual-level analyses consistently indicate that real fish behave as if they interact effectively with the single most influential neighbor. Real fish showed greater directional variability than virtual fish, broader distributions of turning angles, and weaker alignment with the wall when coordination among virtual fish increased. These patterns are consistent with a mechanism in which the identity of the most influential neighbor changes frequently, producing directional fluctuations. Experimental and theoretical studies have further suggested that selective attention and rapid switching between targets are widespread in fish and other animals, and that such switching can enhance responsiveness and behavioral flexibility without requiring simultaneous integration of multiple stimuli [19, 21, 41, 42].

Cross-correlation analyses support this interpretation. The temporal structure of correlations between the real fish and the virtual fish was best reproduced by simulations assuming *k*_R_ = 1, independently of *k*_V_. By contrast, correlations among virtual fish remained strong and symmetric for all values of *k*_V_, reflecting their tighter coupling. This difference suggests that the real fish is coupled to the group through a simpler interaction rule than the one governing the virtual subgroup. Temporal coupling and reciprocity are increasingly recognized as key determinants of coordination, because the timing of responses can shape both spatial organization and information transfer within groups [28].

Taken together, these results support a parsimonious view of social-information integration in schooling fish. Effective coordination does not necessarily require integrating information from many neighbors simultaneously, in contrast to classical phenomenological models of collective motion, which generally assume that individuals interact with multiple neighbors within a metric or topological neighborhood (e.g., [43–46]). Instead, following a single most influential neighbor, whose identity frequently changes over time, appears sufficient to maintain cohesion and alignment in many situations. Computational and robotic studies have reached similar conclusions showing that simple combinations of attraction and alignment, applied selectively to a limited subset of neighbors, can reproduce realistic schooling dynamics and robust collective responses [20, 21, 42].

Such a strategy may offer important advantages. Limiting the number of neighbors reduces sensory and cognitive demands and may increase robustness in noisy environments. At the same time, switching influential neighbors allows individuals to remain responsive to local perturbations and to rapidly propagate directional changes through the group. Theoretical work further suggests that variability or stochasticity in neighbor choice can enhance cohesion and prevent fragmentation, highlighting the adaptive value of flexible interaction rules [42]. This balance between simplicity and flexibility may be a general principle of distributed coordination emerging across taxa and organizational scales whenever information-processing capacity is limited but responsiveness must remain high [14].

More broadly, our study illustrates the power of biohybrid closed-loop systems for studying collective behavior. By combining real animals with controllable virtual agents, it becomes possible to manipulate interaction rules directly and to test hypotheses that are difficult to address in fully natural groups. This hybrid approach represents a shift from validating models solely through statistical agreement toward testing whether model-driven agents elicit the correct behavioral responses in living organisms, providing a stronger and more mechanistic benchmark for theories of collective behavior [27, 30].

Future studies could extend this approach by introducing heterogeneity in interaction rules or sensory delays, and by exploring larger group sizes. Such experiments may help to bridge the gap between individual decision rules and the emergent dynamics of collective motion across biological and artificial systems, and may ultimately contribute to a unified understanding of how distributed sensing, selective attention, and local interactions give rise to robust collective intelligence.

## IV. MATERIALS AND METHODS

### Ethics statement

Experiments were approved by the Animal Experimentation Ethics Committee C2EA-01 of the Toulouse Biology Research Federation and were performed in an approved fish facility (A3155501) under permit APAFIS#27303-2020090219529069 v8 in agreement with the French legislation. All procedures were designed to minimize stress and handling. Fish were transferred from rearing tanks to the experimental setup with minimal manipulation. Each individual was used in only one one-hour experimental session per week. Swimming ability was monitored throughout; fish exhibiting impaired or absent swimming activity were excluded and replaced. No animals were sacrificed during this study.

### Study species

Rummy-nose tetras (*Hemigrammus rhodostomus*) were purchased from Amazonie Labège in Toulouse, France. Fish were kept in 16 L aquariums on a 12:12 hour, dark:light photoperiod, at 27.7^*°*^ C (*±* 0.4^*°*^ C) and were fed *ad libitum* with fish flakes. The average body length of the fish used in these experiments is 3.1 cm.

### Experimental setup

Experiments were carried out using a custom immersive virtual-reality platform, previously developed and validated to investigate interactions between a freely swimming fish and virtual conspecifics under closed-loop conditions [29]. In that earlier methodological work, the behavioral responses elicited by virtual stimuli were shown to be statistically indistinguishable from those observed during interactions with real conspecifics, thereby demonstrating the biological realism and experimental validity of the virtual fish.

The setup consisted of a hemispherical acrylic bowl (diameter 52.2 cm, depth 14.6 cm, filled with 15 L of water) positioned directly above a DLP LED projector that displayed anamorphically rendered virtual fish onto the bowl wall (Figure 1A). The virtual fish are rendered anamorphically so as to preserve correct depth and perspective cues as a function of the real fish’s position and orientation. As a result, apparent variations in the projected size of the virtual fish reflect the geometry of immersive projection rather than changes in the perceived size of the stimulus.

Fish movements were tracked in three dimensions using an Intel RealSense D435 depth camera positioned 47.9 cm above the water surface. Infrared illumination (8 IR lamps, 850 nm) enabled robust tracking under controlled light conditions. A dedicated workstation (Dell Precision 3640 with NVIDIA RTX 3070) processed all tasks in real time, including 3D tracking, trajectory simulation of virtual fish, and rendering.

The system architecture consists of three main software components: a Python-based 3D tracking pipeline, a C++ trajectory simulator, and a rendering engine implemented in Unity. In the present study, the system was used in closed-loop mode: the position and heading of the virtual fish were continuously controlled by a three-dimensional behavioral model parameterized for rummy-nose tetras, exploiting the current position and speed of the real fish. In this model, the interaction forces were specifically tuned for each value of *k*_V_ = 1, 2, and 4 to best reproduce previous experimental results obtained with groups of five real fish. Real and virtual fish positions were exchanged via UDP at 30–90 Hz, ensuring precise and biologically relevant interactions. The same configuration was used across all experimental conditions, which differed only in the number of neighbors influencing the behavior of the virtual fish, in order to quantify the response of a single real fish, while keeping the visual environment and illumination strictly identical throughout the experiments.

### Experimental procedure

Before each experiment, the hemispherical virtual-reality (VR) bowl was filled with water taken from the holding tank. A single real fish was then gently transferred from the holding tank to the hemispherical VR bowl. The virtual-reality system was subsequently activated, and the positions and headings of the virtual fish, controlled in real time by the model, were continuously updated in a closed-loop manner based on the instantaneous positions and velocities of the interacting fish. The real fish then interacted with the virtual conspecifics.

Data acquisition began after a 15 min acclimation period, allowing the real fish to adapt to the experimental environment. Each individual participated in a single continuous 1 h experiment. After the experiment, the fish was returned to the holding tank and was not reused. Water temperature was measured and recorded both before and after each experiment.

During the experiments, the behavior of the virtual fish was determined by varying the number of influential neighbors considered by the model: the single most influential neighbor, the two most influential neighbors, or all four neighbors, corresponding to *k*_V_ = 1, 2, and 4, respectively. Model parameters were adjusted accordingly for each *k*_V_ condition (see Supplementary Table 1 for parameter values). Experiments under different *k*_V_ conditions were conducted in an alternating sequence. For each condition, eight independent trials were performed, yielding a total of 8 h of data per *k*_V_ condition (864, 000 frames, recorded at 30 frames per second).

### Preprocessing of fish trajectories

Before trajectory analysis, experimental data were preprocessed to ensure accuracy and to retain only periods in which fish were actively swimming. Time intervals during which a fish remained immobile for more than 4 s were excluded, as were trajectory segments in which the instantaneous swimming speed exceeded 25 cm/s, or in which a fish moved along a straight trajectory at approximately constant speed for more than 1 s. These empirical thresholds were determined from the statistical properties of spontaneous swimming behavior and were chosen to remove periods of inactivity as well as tracking artefacts, such as abrupt position jumps.

The dataset was further filtered to remove any frames in which the reconstructed fish position lay outside the bowl. To reconstruct continuous temporal trajectories, gaps introduced by this filtering procedure were linearly interpolated when their duration was shorter than 1 s. For longer gaps, reliable reconstruction was not possible, and trajectories were therefore left discontinuous at those points.

The resulting discrete trajectories therefore consisted of multiple consecutive temporal segments. Each segment was subsequently smoothed using a Gaussian kernel convolution to reduce tracking noise while preserving the essential kinematic features of the motion,

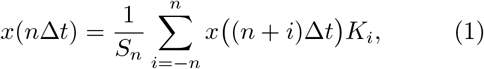

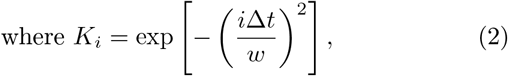

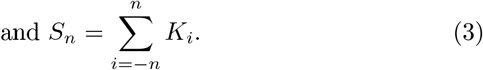

We use *n* = 4*h*/Δ*t*, so that the exponential kernel weight is negligible at *i* = *n*. The kernel width *w* is set to *w* = 0.5 s, which is the typical duration of a kick arising from the burst-and-coast swimming mode of *H. rhodostomus* [9], thus providing minimal smoothing while effectively reducing high-frequency noise.

Although simulation data are intrinsically smoother and subject to far fewer losses than experimental data, both datasets were processed using identical filtering and smoothing procedures to ensure a consistent and unbiased comparison.

## Data analysis

### Quantification of individual and collective behavior in real and bio-hybrid groups of fish

Fish predominantly swim within a horizontal plane at a preferred depth. The experimental tank was modelled as the lower spherical cap of a sphere with radius *R*_0_ = 26.19 cm, filled with 15 L of water, resulting in a central water depth of *h*_water_ = 14.73 cm. The origin of the coordinate system was defined at the water surface directly above the centre of the container. The *𝓏*-axis pointed downward, such that depth values were positive and increased with vertical distance from the surface.

At a given depth *𝓏*, the horizontal cross-section of the tank had a radius *R*_*𝓏*_ given by

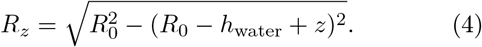

The position and velocity vectors of a fish are 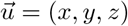 and 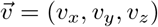 respectively. We define two measures in the horizontal plane: the radial distance to the vertical line defining the *𝓏*-axis, 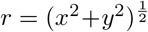, and the horizontal speed 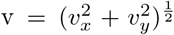 . Then, the distance of the fish to the wall of the bowl on this plane is *r*_w_ = *R*_*𝓏*_ *r*. The fish has an azimuthal (heading) direction angle *ϕ* = atan2(*v*_*y*_, *v*_*x*_), a positional angle *θ* = atan2(*y, x*) on the horizontal plane, and an elevation angle *χ* = atan2(*v*_*𝓏*_/v) relative to the *𝓏*-axis. Accordingly, the angle of incidence of the fish’s trajectory with respect to the wall is denoted as *θ*_w_ = *ϕ θ*.

For two (real or virtual) fish *i* and *j*, the three-dimensional distance between them is 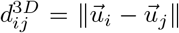, so their horizontal and vertical separations are *d*_*ij*_ 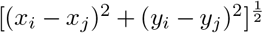 and 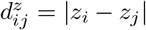 respectively. The difference in heading orientation within the horizontal plane is *ϕ*_*ij*_ = *ϕ*_*j*_ *−ϕ*_*i*_, while the difference in pitch angle along the vertical axis is *𝓏*-axis is *χ*_*ij*_ = *χ*_*j*_ *−χ*_*i*_. From the perspective of fish *i*, the visual angle toward fish *j*, that is, the angular correction required for fish *i* to orient toward fish *j* in the horizontal plane, is defined as *ψ*_*ij*_ = atan2(*y*_*j*_ *−y*_*i*_, *x*_*j*_ *−x*_*i*_) *−ϕ*_*i*_.

The position and velocity vectors of the barycenter of all (real and virtual) individuals are given by

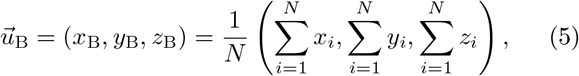

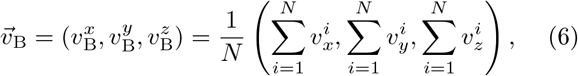

where *N* is the total number of individuals.

In this study, quantities associated with real fish are denoted by the index “R”, whereas those associated with virtual fish are denoted by the index “V”. We further define the barycenter of the virtual fish, whose position and velocity vectors, 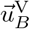 and 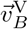 are computed as the averages over the *N −* 1 virtual fish.

The quantification of collective behavior in groups was based on two standard measures: polarization and milling. Because fish predominantly swim within a horizontal plane, only the horizontal components of polarization were considered

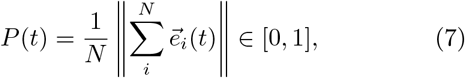

where 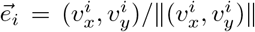 is the unit vector pointing in the heading direction of fish *i*, projected onto the (*x, y*) plane. High values of *P* indicate strong directional alignment among individuals, with *P* = 1 corresponding to perfectly aligned headings.

The milling index *M* quantifies the degree of collective rotational motion around the barycenter of the group. Variables expressed in the reference frame of the barycenter *B* are denoted with an overbar. For instance, 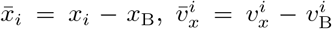 denote the position and velocity components of fish *i* relative to the barycenter. The heading angle in the barycentric frame is defined as 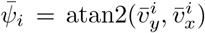, while the angular position of fish *i* relative to the barycenter is 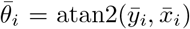. The angular difference 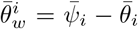 therefore characterizes the tangential component of motion. The milling index is then defined as

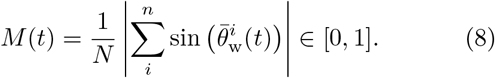

Larger values of *M* indicate stronger collective rotation of the group around its barycenter.

### Statistical metric for probability distribution comparison

To quantitatively compare probability distribution functions (PDFs) obtained under different experimental conditions, we used the Hellinger distance as a measure of similarity [47, 48]. Given two normalized PDFs, *F*(*x*) and *G*(*x*), corresponding to the same observable, the square of the Hellinger distance is defined as

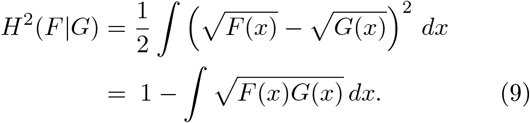

This metric ranges from 0 to 1, where *H*(*F* ∣ *G*) = 0 if and only if the two distributions are identical, and values approaching 1 indicate distributions with almost non-overlapping support. The second form of the equation provides a straightforward interpretation: it measures the deviation from unity of the scalar product between the square-rooted distributions, viewed as unit vectors in Euclidean space. In practice, a small Hellinger distance (*H <* 0.2) indicates a high similarity between the distributions, while values greater than 0.2 indicate a meaningful dissimilarity (note that in Supplementary Tables 2 and 5, all values have been scaled (×10) to facilitate comparison). This approach provides a robust and interpretable quantification of differences between behavioral distributions and has been successfully applied to assess the agreement between empirical fish data and model simulations in collective behavior studies.

### Correlation analyses

To quantify the similarity of dynamical behaviors, we computed the normalized cross-correlation function of velocity vectors. For two individuals *i* and *j* with velocity vectors 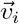 and 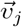, the cross-correlation at time lag *τ* was defined as

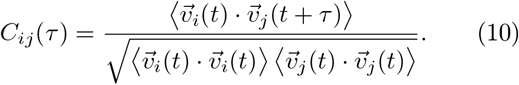

This measure was used both to quantify coordination between the real fish and its nearest virtual neighbor (*i* = R, *j* = V) and to characterize coordination between pairs of distinct virtual fish (*i* = V_*m*_, *j* = V_*n*_, with *m ≠ n*).

### 3D self-propelled particle model for fish swimming in a bowl

The temporal variation of the position of fish *i* is given by the evolution of its three-dimensional acceleration vector 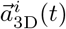,

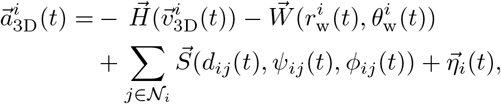

where 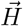 represents the hydrodynamic friction and speed adaptation to neighbors, which depends on the instantaneous velocity of the individual; 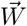 accounts for the horizontal wall effects, which depend on the distance and angle of incidence to the wall; 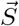 describes social interactions with neighboring fish, determined by the pairwise relative state between the focal fish *i* and each interacting neighbor *j* given by the distance, the angle of perception, and the heading difference between the focal fish *i* and the neighbor *j* including their mutual distance, viewing angle, and heading difference. The term 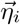 is a noise term accounting for the intrinsic behavioral fluctuations of fish *i*.

Decomposing the acceleration into its components, the horizontal acceleration comprises a longitudinal component associated with changes in speed along the direction of planar motion, defined by the unit vector 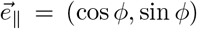, and a lateral component associated with turning within the horizontal plane, oriented perpendicular to the direction of motion, defined by 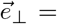 (*−*sin *ϕ*, cos *ϕ*). In addition, a vertical acceleration component acts along the vertical axis, in the direction of the unit vector 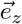.

In the horizontal plane, the longitudinal acceleration is governed by hydrodynamic effects, repulsion from the nearest wall, and attraction toward other individuals. The lateral acceleration is determined by wall-induced repulsion, the tendency of individuals to follow the wall, and social interactions with conspecifics, including attraction and alignment. Vertical acceleration reflects the tendency of fish to swim at a preferred depth. When conspecifics occupy different depths, they exert an attractive influence that drives the focal individual toward their depth. Accordingly, vertical acceleration is governed by the relaxation of vertical velocity toward zero, the restoring force toward a preferred depth, and the vertical attraction exerted by individuals located at different depths.

This leads us to the following system of equations for fish *i*,

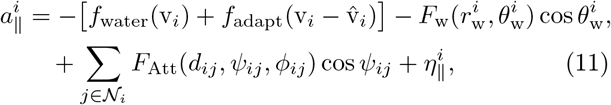

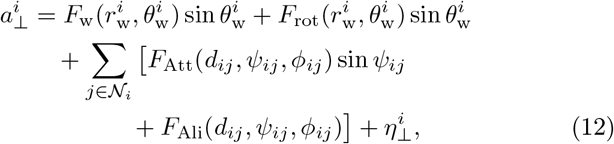

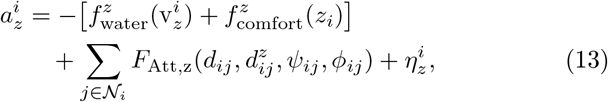

where 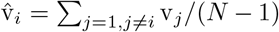, and *𝒩*_*i*_ denotes the set of the neighbors interacting with fish *i*.

The *influence ℐij* that fish *j* exerts on fish *i* is defined as the instantaneous contribution of fish *j* to the change in velocity of fish *i*. It is quantified as

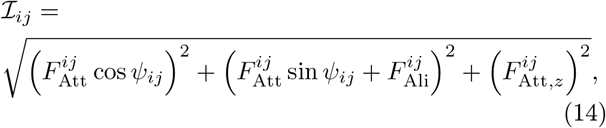

where the dependence of the forces on *t, d*_*ij*_, *ψ*_*ij*_, *ϕ*_*ij*_ (and 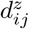 for *F*_Att, *𝓏*_) has been omitted for clarity.

The strategy by which social information is processed is determined by the number *k* of neighbors included in the interaction set 𝒩_*i*_, ranked according to their instantaneous influence on the focal fish *i*. Neglecting less influential neighbors reduces the amount of information that must be processed by the fish’s nervous system and limits potential cognitive overload. We considered three cases: *k* = 1, 2, and 4. For interactions involving virtual fish—both in experiments and simulations, including interactions among virtual fish and between virtual and real fish—we denote this parameter as *k*_V_. For interactions involving simulated real fish interacting with virtual fish in simulations, we use the notation *k*_R_. Because influence is defined as an instantaneous quantity and relative spatial configurations continuously evolve, the identity of the *k* most influential neighbors may change from one time step to the next.

We also assume that the multi-variable functions can be decomposed into the product of single-variable functions, e.g., *F*_Att_(*d*_*ij*_, *ψ*_*ij*_, *ϕ*_*ij*_) = *F*_Att_(*d*_*ij*_) *G*_Att_(*ψ*_*ij*_) *h*_Att_(*ϕ*_*ij*_) and 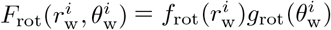, and similarly for *F*_w_, *F*_Ali_, and *F*_Att,z_; see the analytical expressions of the functions in the Supplementary Material.

Each fish has its own autocorrelated noises 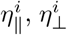, and 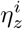 in the parallel, perpendicular, and vertical directions, respectively, which evolve according to a discrete Ornstein–Uhlenbeck process, expressed as

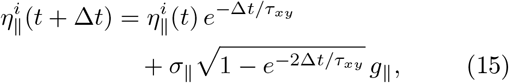

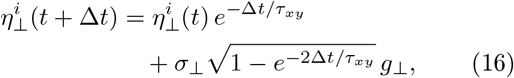

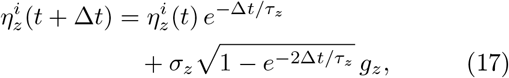

where *g*_∥_, *g*_*⊥*_, and g_*𝓏*_ are independent standard normal deviates. These random deviates are generated using the Box–Muller transform,

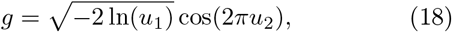

with *u*_1_, *u*_2_ *∼U*(0, 1). Here, *σ*_*∥*_, *σ*_*⊥*_, and *σ*_*z*_ denote the noise amplitudes, while *τ*_*xy*_ and *τ*_*z*_ represent the correlation time, which are not necessarily the same in the horizontal and vertical components. The use of finite correlation times is consistent with the burst-and-coast swimming mode of fish, in which brief acceleration bursts (typically lasting about 0.1 s) alternate with passive gliding phases (typically about 0.5 s), during which the fish exhibits only minimal changes in heading.

It may happen that, at some iteration, the predicted head position is outside the water volume. Such moves are therefore rejected. The coordinates (*x, y, 𝓏*) denote the center of mass of the fish. The position of its head is given by *x*_*h*_ = *x* + *l*_*h*_ cos *ϕ* cos *χ, y*_*h*_ = *y* + *l*_*h*_ sin *ϕ* cos *χ, 𝓏*_*h*_ = *𝓏* + *l*_*h*_ sin *χ*, where *h* is the head-center distance. When *R*_w_(*x*_*h*_(*t* + Δ*t*), *y*_*h*_(*t* + Δ*t*), *𝓏*_*h*_(*t* + Δ*t*)) *<* 0, the fish is returned to its previous position (*x*(*t*), *y*(*t*), *𝓏* (*t*)), its orientation is adjusted nearly tangentially to the wall, and the sign of the perpendicular noise is adjusted to point in the same direction:

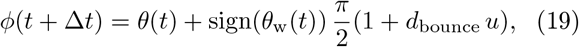

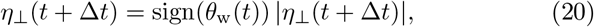

where *d*_bounce_ = 0.25 is a small random angular correction and *u ∼U*(0, 1). This ensures that the fish head is nearly aligned with the wall, *θ*_w_(*t* + Δ*t*) *≈ ±π/*2.

In rare cases where the predicted head position lies above the water surface or below the bottom of the bowl, a corrective step is applied in which the depth is reset and the vertical velocity is reversed, such that: *𝓏* (*t* + Δ*t*) = *𝓏* (*t*), *v*_*𝓏*_ (*t* + Δ*t*) = *−v*_*𝓏*_ (*t*).

Following this correction, numerical integration proceeds normally using the updated state variables. Additional details on the numerical integration scheme are provided in the Supplementary Material.

### Determination of model parameters

Model parameters were fitted using two-dimensional trajectory data obtained from recent experiments conducted on groups of five real fish [38]. In these experiments, collective trajectories were recorded in a circular arena of radius *R* = 250 mm, matching the diameter of the virtual-reality setup, with a water depth of 7 cm and a controlled illumination level of 50 lux, comparable to that measured in the virtual-reality system. Because fish typically swam within a vertical range of 2 to 6 cm, the effective swimming layer and visual conditions were similar in both environments, resulting in closely comparable collective motion patterns. The fitting procedure was designed to ensure that simulated trajectories quantitatively reproduced the empirical distributions of a set of five individual- and collective-level behavioral metrics measured in the corresponding experiments. These metrics include the probability density functions of swimming speed v, the distance and angle of incidence to the wall *r*_w_ and *θ*_w_, the distance between fish *d*, the viewing *ψ*, and the alignment *ϕ*.

Four key parameters were tuned by introducing a specific coefficient for each interaction term: *i*) the individual swimming speed, *ii*) the strength of wall repulsion, *iii*) the inter-individual attraction strength, and *iv*) the alignment strength. A distinct set of parameter values was obtained for each value of *k* = 1, 2, and 4 (Supplementary Table 1).

As an illustration of the fitting results, the experimentally measured mean distance of individuals from the tank wall, *r*_w_ *≈*6.8 cm, lies within the range of mean values generated by the model for *k*_V_ = 1, 2, and 4, which spans from *r*_w_ *≈*8.6 to 5.5 cm. Similar qualitative agreement was observed for the remaining five metrics.

## DATA AVAILABILITY

The datasets generated and/or analyzed during the current study are available at: https://figshare.com/s/65f5950fae41ee24ecbf.

All codes that support the findings of this study are available on figshare with the identifier: https://figshare.com/s/bcc1919ab042c998fc5c.

## ACKNOWLEDGMENTS

This work was funded by Agence Nationale de la Recherche (ANR-20-CE45-0006). ZK was supported by a grant from the China Scholarship Council (CSC N^*°*^202406040167). RE was partially supported by the Spanish AEI grant PID2020-115088RB-I00. The founders had no role in study design, data collection and analysis, decision to publish, or preparation of the manuscript.

