## Supplemental Data 1 for "Decoding social integration in schooling fish using real–virtual interactions"

*guy.theraulaz@univ-tlse3.f

This PDF file includes:

- Supplemental text
- Supplemental Figures S1 to S10
- Supplemental Tables S1 to S5
- Supplemental Movies S1 to S7

### SUPPLEMENTAL TEXT

#### *Numerical integration*

In the numerical simulations, the model is formulated in Cartesian coordinates, which can be recovered with $a_x^i = a_\parallel^i \cos\phi_i - a_\perp^i \sin\phi_i$ and $a_y^i = a_\parallel^i \sin\phi_i + a_\perp^i \cos\phi_i$, together with the identities

$$\begin{aligned}
\cos\theta_{\rm w}^i \cos\phi_i + \sin\theta_{\rm w}^i \sin\phi_i &= \cos(\theta_{\rm w}^i - \phi_i) = \cos(-\theta_i) = \cos\theta_i,\\
\cos\theta_{\rm w}^i \sin\phi_i - \sin\theta_{\rm w}^i \cos\phi_i &= \sin(\phi_i - \theta_{\rm w}^i) = \sin\theta_i,\\
\cos\psi_{ij} \cos\phi_i - \sin\psi_{ij} \sin\phi_i &= \cos(\psi_{ij} + \phi_i) = \cos\theta_{ij},\\
\cos\psi_{ij} \sin\phi_i + \sin\psi_{ij} \cos\phi_i &= \sin(\psi_{ij} + \phi_i) = \sin\theta_{ij}.
\end{aligned}$$

Then, the equations are (the one for the vertical component $a_z^i$ is unchanged),

$$\begin{aligned}
a_x^i = &-[f_{\rm water}({\rm v}_i) + f_{\rm adapt}({\rm v}_i - \hat{\rm v}_i)]\cos\phi_i - F_{\rm w}(r_{\rm w}^i, \theta_{\rm w}^i)\cos\theta_i - F_{\rm rot}(r_{\rm w}^i, \theta_{\rm w}^i)\sin\theta_{\rm w}^i \sin\phi_i\\
&+ \sum_{j\in\mathcal{N}_i} F_{\rm Att}(d_{ij}, \psi_{ij}, \phi_{ij})\cos\theta_{ij} - \sum_{j\in\mathcal{N}_i} F_{\rm Ali}(d_{ij}, \psi_{ij}, \phi_{ij})\sin\phi_i + \eta_\parallel^i \cos\phi_i - \eta_\perp^i \sin\phi_i,\\
a_y^i = &-[f_{\rm water}({\rm v}_i) + f_{\rm adapt}({\rm v}_i - \hat{\rm v}_i)]\sin\phi_i + F_{\rm w}(r_{\rm w}^i, \theta_{\rm w}^i)\sin\theta_i + F_{\rm rot}(r_{\rm w}^i, \theta_{\rm w}^i)\sin\theta_{\rm w}^i \cos\phi_i\\
&+ \sum_{j\in\mathcal{N}_i} F_{\rm Att}(d_{ij}, \psi_{ij}, \phi_{ij})\sin\theta_{ij} + \sum_{j\in\mathcal{N}_i} F_{\rm Ali}(d_{ij}, \psi_{ij}, \phi_{ij})\cos\phi_i + \eta_\parallel^i \sin\phi_i + \eta_\perp^i \cos\phi_i,
\end{aligned}$$

and the influence is

$$\mathcal{I}_{ij} = \sqrt{\left(F_{\rm Att}^{ij}\cos\theta_{ij} - F_{\rm Ali}^{ij}\sin\phi_i\right)^2 + \left(F_{\rm Att}^{ij}\sin\theta_{ij} + F_{\rm Ali}^{ij}\cos\phi_i\right)^2 + \left(F_{{\rm Att},z}^{ij}\right)^2}. \tag{S1}$$

#### *Analytical expressions of the extracted interaction functions*

The decomposition of the equations of motion allows us to incorporate the fundamental symmetry constraints that the functional form of the angular terms must verify to reproduce the directional reactions of fish. These constraints preserve the right-left symmetry, which was shown to consistently hold for the considered fish species.

For instance, the rotational force $F_{\rm rot}$ must include a factor $\sin(\theta_{\rm w})$ ensuring that a fish approaching the wall with $\theta_{\rm w} > 0$ (wall to its right) is pushed to turn left with the same intensity as it is pushed to turn right when it approaches the wall with $\theta_{\rm w} < 0$ (wall to its left). Similarly, the alignment force $F_{\rm Ali}$ must include a factor $\sin(\Delta\phi)$ favoring a left turn when the neighbor is misaligned to the left ($\Delta\phi > 0$), thus increasing the perpendicular acceleration, while if the neighbor is misaligned to the right ($\Delta\phi < 0$), the contribution must be negative to induce a right turn. These constraints preserve the right-left symmetry, which was shown to consistently hold for the considered fish species.

We assume that, once decomposed along the parallel and perpendicular components of the acceleration, the forces can be separated as products of single-variable functions, so the forces accounting for the effect of the wall are written as,

$$F_{\rm w}(r_{\rm w}, \theta_{\rm w}) = f_{\rm w}(r_{\rm w})\, g_{\rm w}(\theta_{\rm w}), \tag{S2}$$
$$F_{\rm rot}(r_{\rm w}, \theta_{\rm w}) = f_{\rm rot}(r_{\rm w})\, g_{\rm rot}(\theta_{\rm w})\sin(\theta_{\rm w}), \tag{S3}$$

while those representing social interactions are:

$$F_{\rm Att}(d, \psi, \Delta\phi) = f_{\rm Att}(d)\, g_{\rm Att}(\psi)\, h_{\rm Att}(\Delta\phi), \tag{S4}$$
$$F_{\rm Ali}(d, \psi, \Delta\phi) = f_{\rm Ali}(d)\, g_{\rm Ali}(\psi)\, h_{\rm Ali}(\Delta\phi)\sin(\Delta\phi), \tag{S5}$$
$$F_{\rm Att}^z(d, d_z, \psi, \Delta\phi) = f_{\rm Att}^z(d)\, k_{\rm Att}^z(d_z)\, g_{\rm Att}^z(\psi)\, h_{\rm Att}^z(\Delta\phi), \tag{S6}$$

where functions denoted by $g$ and $h$ are even.

*Horizontal components:*

$$f_{\mathrm{water}}(\mathrm{v}) = C_{\mathrm{v}}^{(0)} + C_{\mathrm{v}}^{(1)}(\mathrm{v}-\mathrm{v}_0) + C_{\mathrm{v}}^{(2)}(\mathrm{v}-\mathrm{v}_0)^2 + C_{\mathrm{v}}^{(3)}(\mathrm{v}-\mathrm{v}_0)^3, \tag{S7}$$

$$f_{\mathrm{adapt}}(\mathrm{v}) = C_{\mathrm{v}}^{(4)}\mathrm{v} + C_{\mathrm{v}}^{(5)}\mathrm{v}^2 + C_{\mathrm{v}}^{(6)}\mathrm{v}^3, \tag{S8}$$

$$f_{\mathrm{w}}(r_{\mathrm{w}}) = C_{f_{\mathrm{w}}}^{(1)} \exp\left[-\frac{r_{\mathrm{w}}}{C_{f_{\mathrm{w}}}^{(2)}} - \left(\frac{r_{\mathrm{w}}}{C_{f_{\mathrm{w}}}^{(3)}}\right)^2\right] - C_{f_{\mathrm{w}}}^{(4)}, \tag{S9}$$

$$f_{\mathrm{rot}}(r_{\mathrm{w}}) = C_{f_{\mathrm{rot}}}^{(1)} \exp\left[-\frac{r_{\mathrm{w}}}{C_{f_{\mathrm{rot}}}^{(2)}} - \left(\frac{r_{\mathrm{w}}}{C_{f_{\mathrm{rot}}}^{(3)}}\right)^2\right] - C_{f_{\mathrm{rot}}}^{(4)}, \tag{S10}$$

$$g_{\mathrm{w}}(\theta_{\mathrm{w}}) = 1 + \sum_{m=1}^{6} C_{\mathrm{w}}^{(m)} \cos(m\theta_{\mathrm{w}}), \quad g_{\mathrm{rot}}(\theta_{\mathrm{w}}) = 1 + \sum_{m=1}^{6} C_{\mathrm{rot}}^{(m)} \cos(m\theta_{\mathrm{w}}), \tag{S11}$$

$$f_{\mathrm{Att}}(d) = \frac{d - C_{f_{\mathrm{Att}}}^{(3)}}{C_{f_{\mathrm{Att}}}^{(1)}} \times \left[1 + \left(\frac{d}{C_{f_{\mathrm{Att}}}^{(2)}}\right)^2\right]^{-C_{f_{\mathrm{Att}}}^{(4)}}, \tag{S12}$$

$$\text{If } d < d_{\mathrm{c}}^{\mathrm{Att}},\ f_{\mathrm{Att}}(d) = f_{\mathrm{Att}}(d) - \left[\left(\frac{d_{\mathrm{c}}^{\mathrm{Att}}}{d}\right)^2 - 1\right] \frac{d_{\mathrm{c}}^{\mathrm{Att}}}{C_{f_{\mathrm{Att}}}^{(1)}}; \quad \text{If } f_{\mathrm{Att}} < f_{\mathrm{Att}}^{\min},\ f_{\mathrm{Att}} = f_{\mathrm{Att}}^{\min}, \tag{S13}$$

$$f_{\mathrm{Ali}}(d) = \frac{d - C_{f_{\mathrm{Ali}}}^{(3)}}{C_{f_{\mathrm{Ali}}}^{(1)}} \times \left[1 + \left(\frac{d}{C_{f_{\mathrm{Ali}}}^{(2)}}\right)^2\right]^{-C_{f_{\mathrm{Ali}}}^{(4)}}, \tag{S14}$$

$$E_{\mathrm{Att}}(\psi) = 1 + \sum_{m=1}^{6} C_{\mathrm{Att}}^{(m)} \cos(m\psi), \quad G_{\mathrm{Att}}(\Delta\phi) = 1 + \sum_{m=1}^{6} D_{\mathrm{Att}}^{(m)} \cos(m\Delta\phi), \tag{S15}$$

$$E_{\mathrm{Ali}}(\psi) = 1 + \sum_{m=1}^{6} C_{\mathrm{Ali}}^{(m)} \cos(m\psi), \quad G_{\mathrm{Ali}}(\Delta\phi) = 1 + \sum_{m=1}^{6} D_{\mathrm{Ali}}^{(m)} \cos(m\Delta\phi). \tag{S16}$$

*Vertical components:*

$$f_z(z) = C_{f_z}^{(0)} + C_{f_z}^{(1)}(z-z_0) + C_{f_z}^{(2)}(z-z_0)^2 + C_{f_z}^{(3)}(z-z_0)^3 + C_{f_z}^{(4)}(z-z_0)^4, \tag{S17}$$

$$f_{\mathrm{water}}^{z}(\mathrm{v}_z) = C_{\mathrm{v}_z}^{(1)}\mathrm{v}_z + C_{\mathrm{v}_z}^{(2)}\mathrm{v}_z^2 + C_{\mathrm{v}_z}^{(3)}\mathrm{v}_z^3, \tag{S18}$$

$$f_{\mathrm{Att}}^{z}(d_z) = C_{f_{\mathrm{Att},z}}^{(1)} d_z + C_{f_{\mathrm{Att},z}}^{(2)} d_z^2 + C_{f_{\mathrm{Att},z}}^{(3)} d_z^3 + C_{f_{\mathrm{Att},z}}^{(4)} d_z^4, \tag{S19}$$

$$f_{\mathrm{Att},z,d}(d) = C_{f_{\mathrm{Att},z,d}}^{(1)} \left[1 + \left(\frac{d}{C_{f_{\mathrm{Att},z,d}}^{(2)}}\right)^{C_{f_{\mathrm{Att},z,d}}^{(3)}}\right]^{-1}, \tag{S20}$$

$$E_{\mathrm{Att}}^{z}(\psi) = 1 + \sum_{m=1}^{6} C_{\mathrm{Att},z}^{(m)} \cos(m\psi), \quad G_{\mathrm{Att}}^{z}(\Delta\phi) = 1 + \sum_{m=1}^{6} D_{\mathrm{Att},z}^{(m)} \cos(m\Delta\phi). \tag{S21}$$

### SUPPLEMENTAL FIGURES

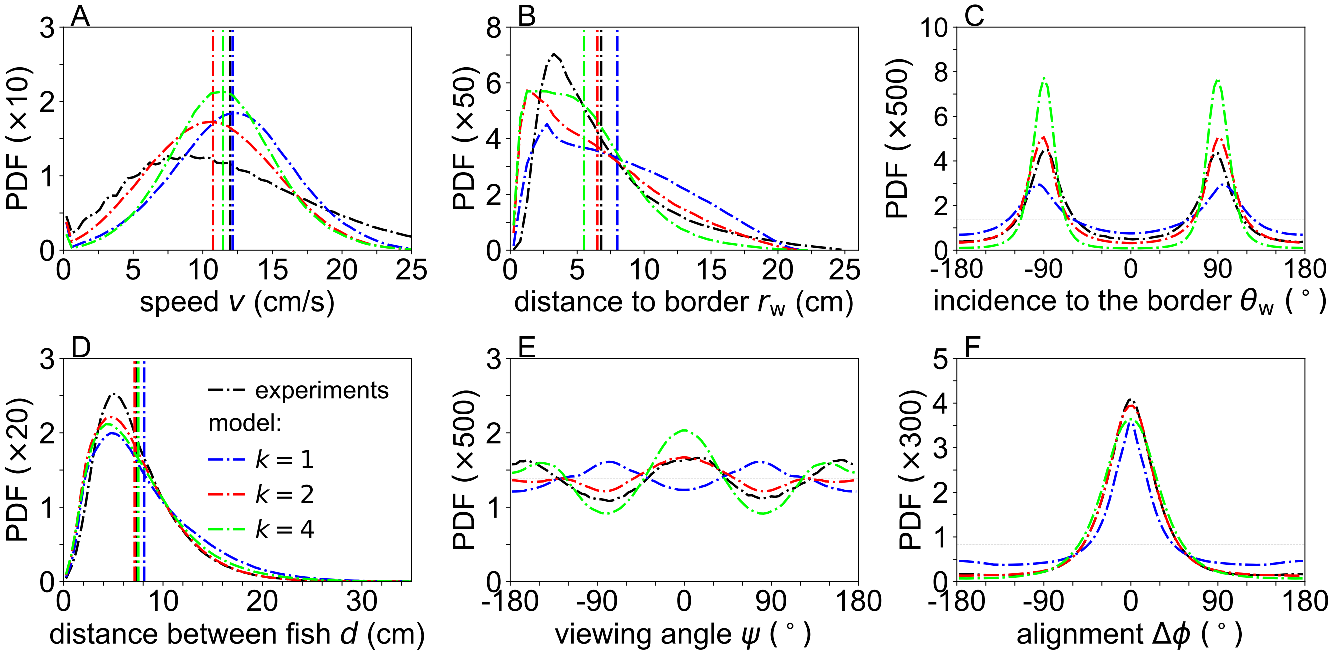


FIG. S1. **Calibration of the 3D behavioral model from 2D trajectory data in groups of five fish**. Distributions of individual kinematic variables and pairwise interaction metrics measured from two-dimensional trajectories recorded in the experiments of Xue et al. (2023), used to estimate correction coefficients for the four interaction forces of the 3D model (individual speed control, wall effects, attraction, and alignment). Probability density functions (PDFs) of (**A**) individual swimming speed $v$, (**B**) distance to the wall $r_w$, (**C**) angle of incidence to the wall $\theta_w$, (**D**) inter-individual distance $d$, (**E**) viewing angle $\psi$, and (**F**) heading-alignment angle $\Delta\phi$. Black lines correspond to experimental data extracted from 2D trajectories. Colored lines show model simulations in which all five individuals follow the same social interaction strategy, for $k = 1$, 2, and 4.

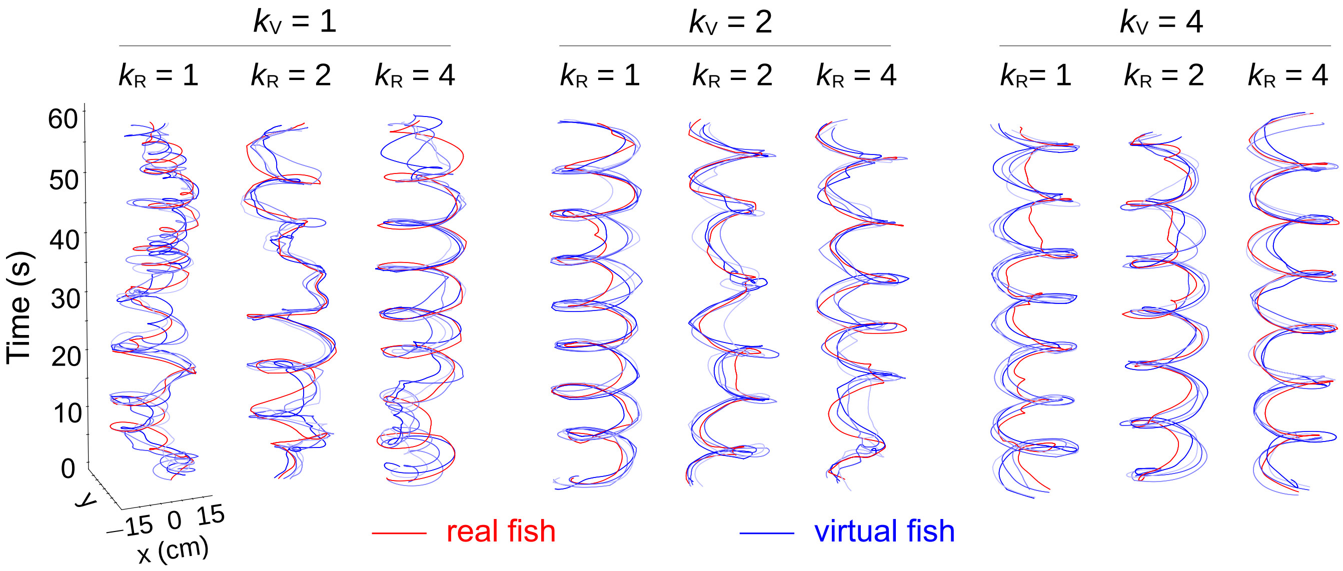


FIG. S2. **Simulated trajectories under different social-information integration strategies.** Representative trajectories of the simulated real fish (red lines) and four virtual conspecifics (blue shades) obtained from nine numerical simulations of the model. Columns correspond to different numbers of influential neighbors considered by the virtual fish: $k_V = 1$, $k_V = 2$ and $k_V = 4$. Within each condition, simulations were performed for three values of the number of influential neighbors considered by the agent representing the real fish, $k_R = 1$, 2, and 4.

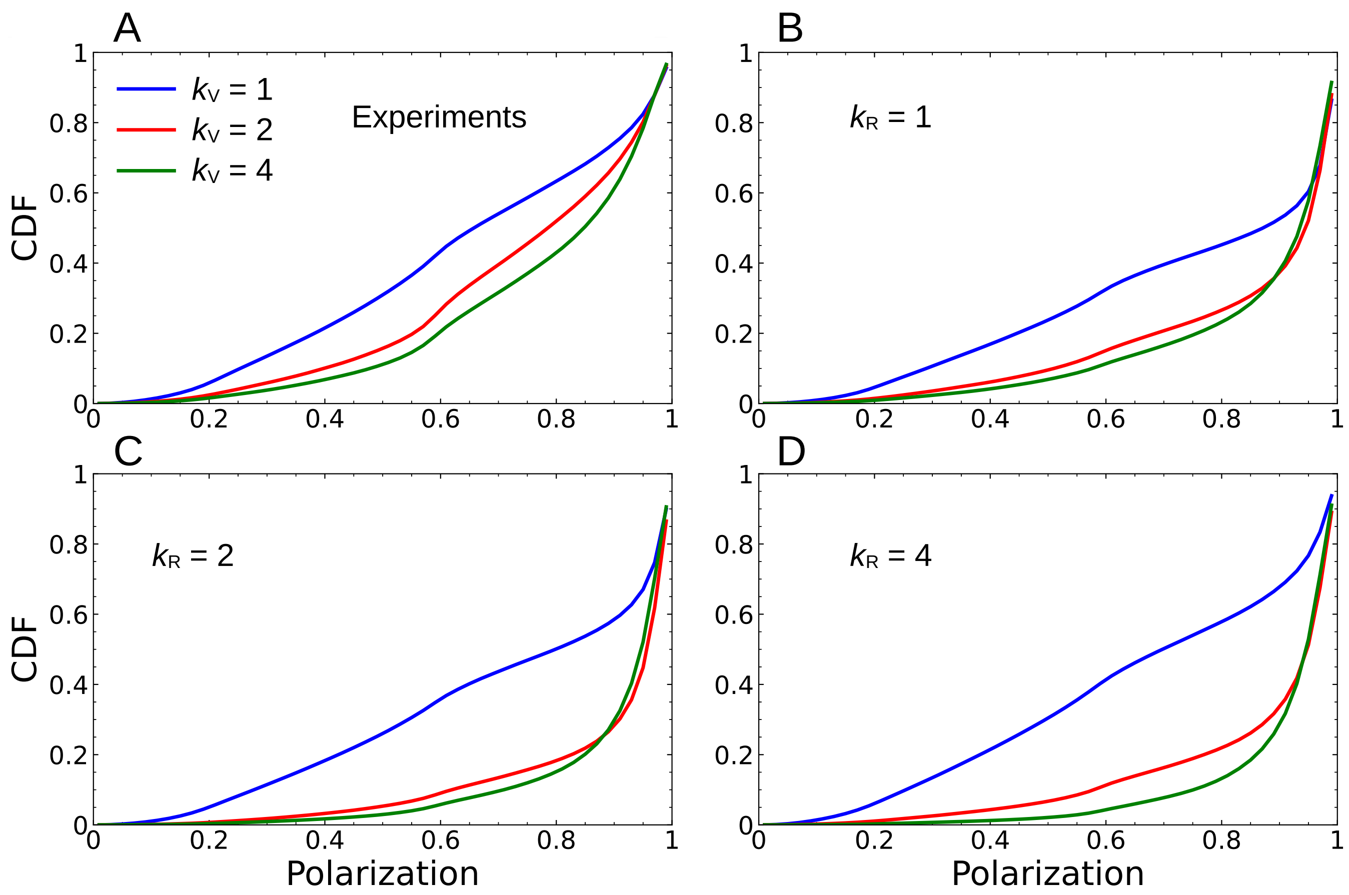


FIG. S3. **Cumulative density function (CDF) of group polarization of the five fish during in experiments and simulations for different social interaction strategies of the virtual fish, $\mathbf{k_V = 1}$ (blue), $\mathbf{k_V = 2}$ (red), and $\mathbf{k_V = 4}$ (green).** (**A**) Experiments. (**B**, **C**, **D**) Model simulations for $k_R = 1$, 2, and 4 respectively.

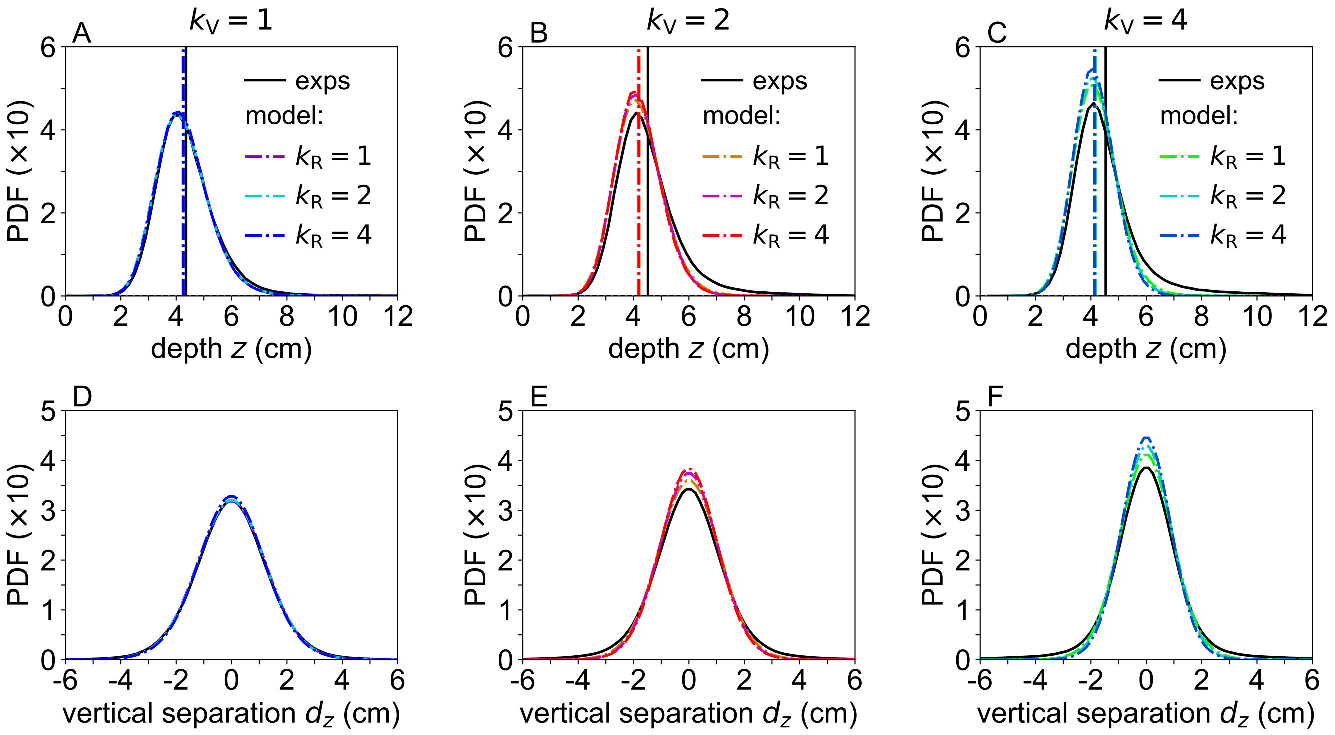


FIG. S4. **Swimming depth and vertical spacing in bio-hybrid groups.** Quantification of vertical motion in bio-hybrid groups composed of one real fish and four virtual conspecifics, averaged over all five individuals. Probability density functions (PDFs) of (**A**–**C**) swimming depth $z$ and (**D**–**F**) vertical separation $d_z$, defined as the mean vertical distance over all pairwise combinations of individuals in the group, for $k_V = 1$ (**A**, **D**), $k_V = 2$ (**B**, **E**), and $k_V = 4$ (**C**, **F**). Black solid lines correspond to experimental data. Colored dashed lines represent model simulations for each value of $k_V$ and for the three social interaction strategies of the real fish, $k_R = 1$, 2, and 4. Vertical lines indicate the mean value of the corresponding PDF in matching colors.

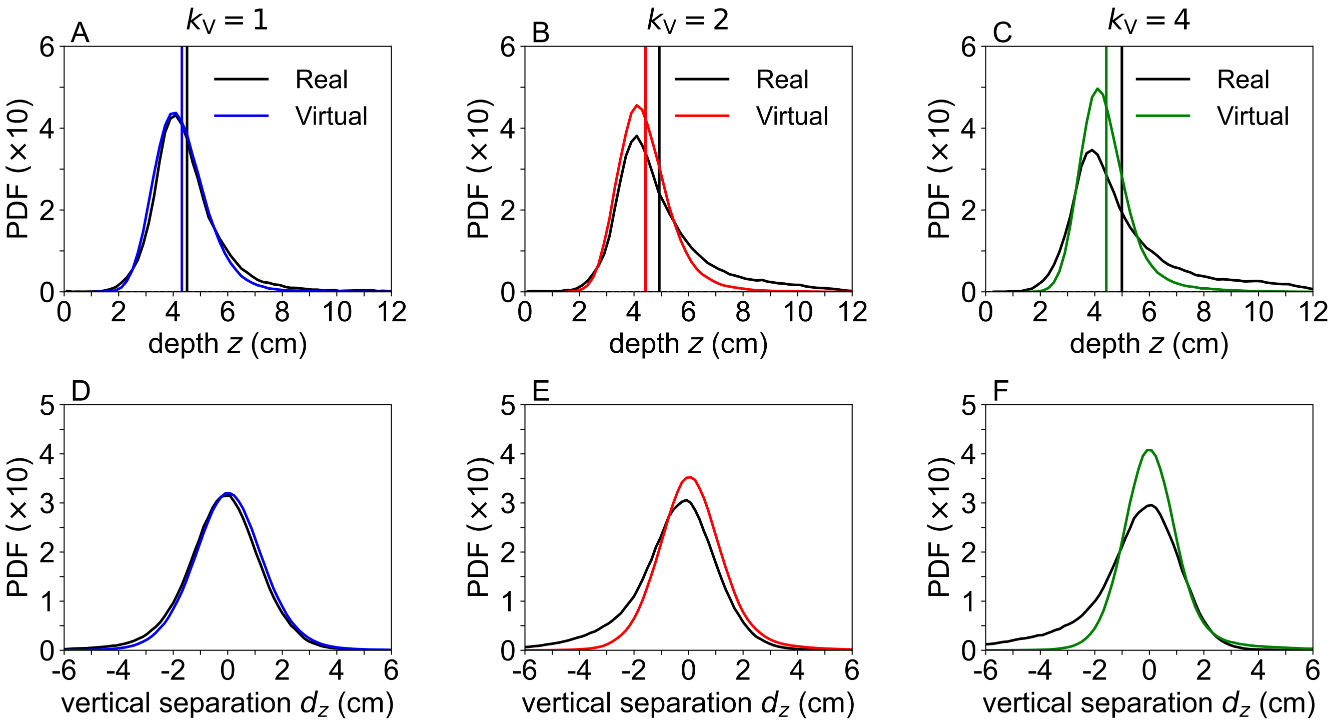


FIG. S5. **Swimming depth and vertical spacing in bio-hybrid groups under different social interaction strategies.** Quantification of the vertical motion of the real fish (black lines) compared with the averaged vertical motion of the four virtual conspecifics (colored lines) during the experiments, for different social interaction strategies of the virtual fish: $k_V = 1$ (**A**, **D**), $k_V = 2$ (**B**, **E**), and $k_V = 4$ (**C**, **F**). Probability density functions (PDFs) of (**A**–**C**) swimming depth $z$ and (**D**–**F**) vertical separation $d_z$. In panels (**D**–**F**), the black curves correspond to the vertical separation averaged over all real–virtual fish pairs, whereas the colored curves correspond to the vertical separation averaged over all pairs involving each virtual fish and the other group members.Vertical lines indicate the mean value of the corresponding PDF in matching colors.

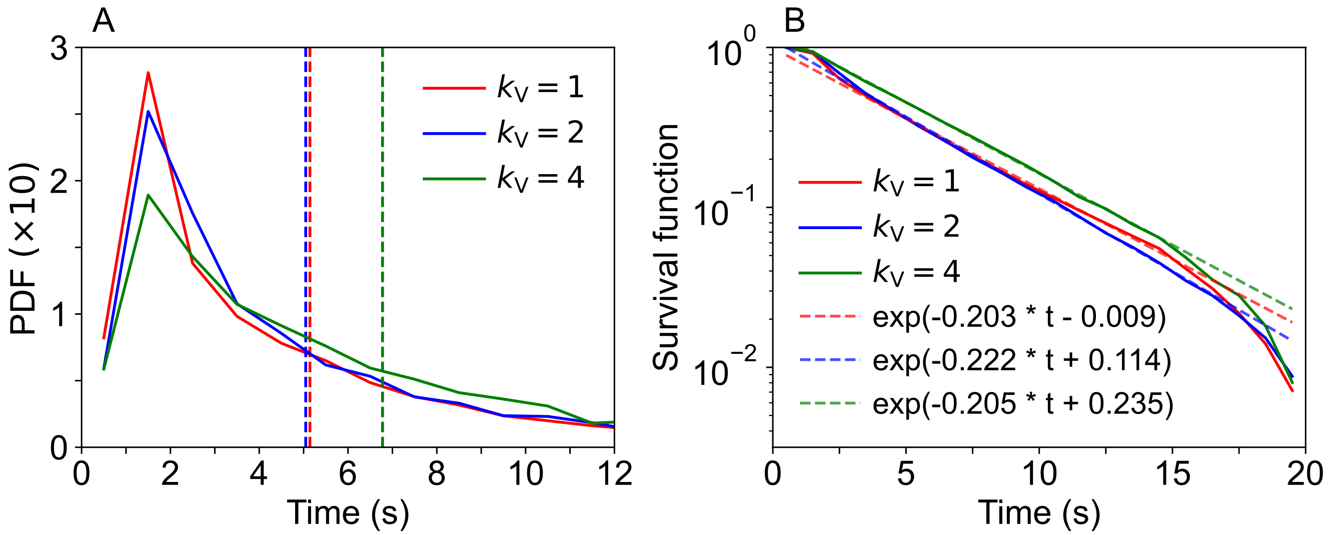


FIG. S6. **U-turn dynamics of the real fish under different levels of group coordination.** Time intervals between successive U-turns of the real fish measured in the experiments for the three social interaction strategies of the virtual fish: $k_V = 1$ (red), $k_V = 2$ (blue), and $k_V = 4$ (green). (**A**) Probability density functions (PDFs) of the time interval between two consecutive U-turns, defined as heading changes exceeding $100^\circ$ within 0.5 s. (**B**) Corresponding survival curves; straight lines indicate exponential fits.

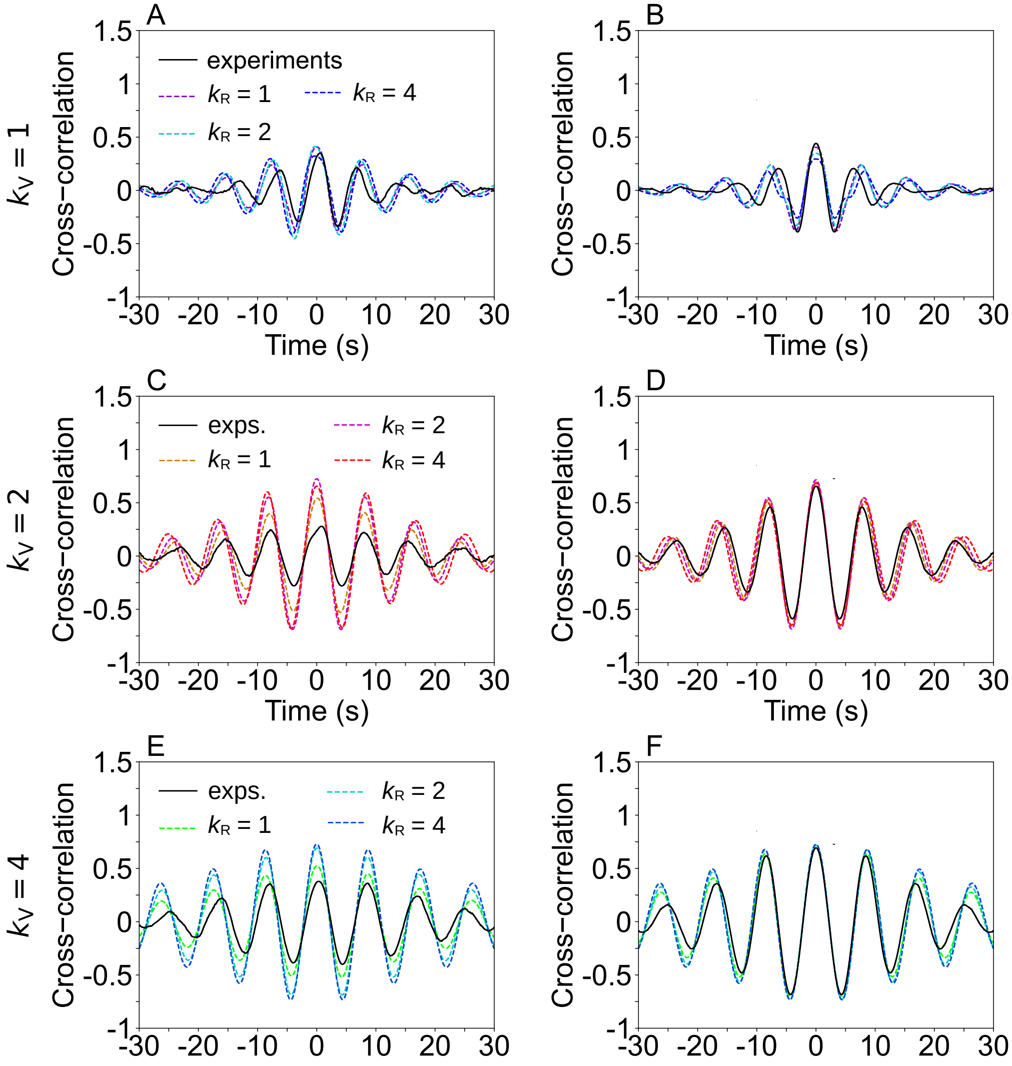


FIG. S7. **Velocity cross-correlations reveal interaction dynamics in bio-hybrid groups.** Cross-correlation functions of velocity vectors between (**A, C, E**) the real fish and a virtual conspecific, and (**B, D, F**) between two distinct virtual fish (cross-correlations only, excluding auto-correlations), for different social interaction strategies of the virtual fish: $k_V = 1$ (first row), $k_V = 2$ (second row), and $k_V = 4$ (third row). Black solid lines correspond to experimental data. Colored lines represent model simulations for the three social interaction strategies of the real fish, $k_R = 1$, 2, and 4.

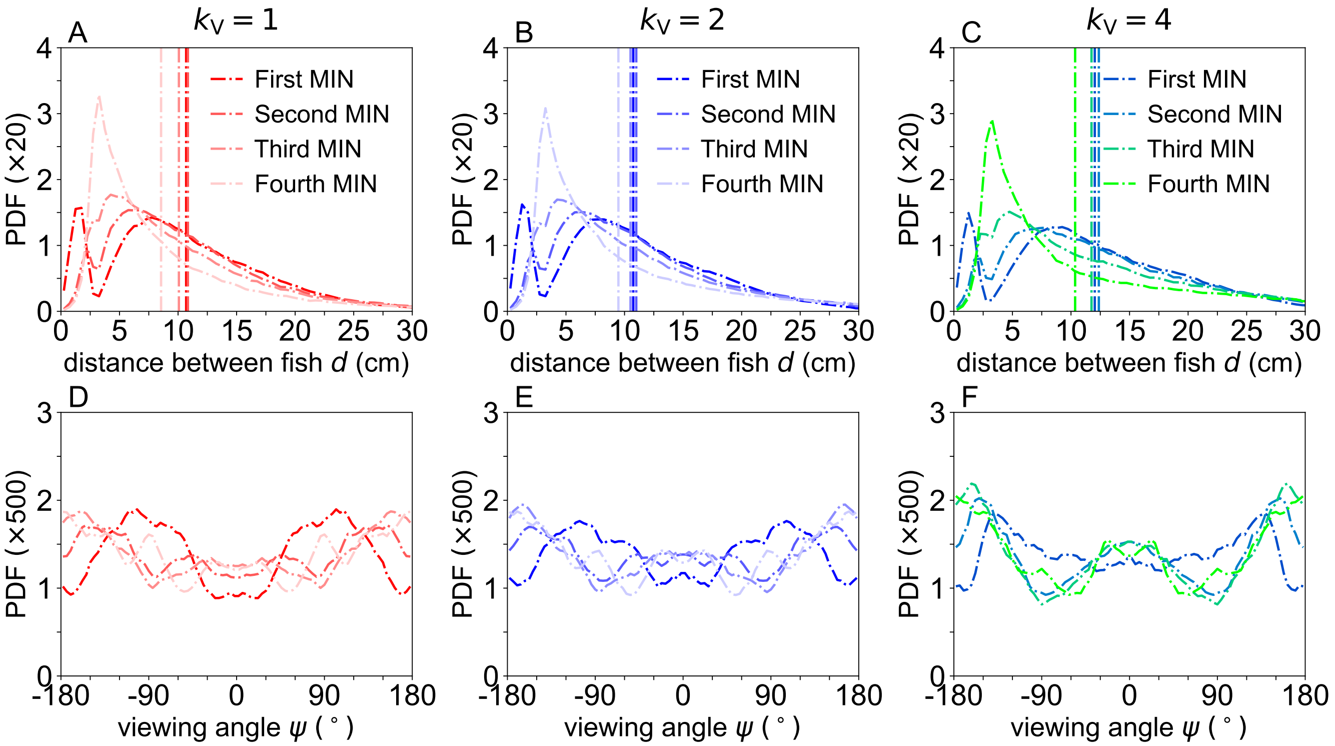


FIG. S8. **Spatial distribution of influential neighbors of the real fish ranked by influence.** Relative positions of the most influential neighbors (MIN) of the real fish measured in the experiments for the three social interaction strategies of the virtual fish, $k_V = 1$, 2, and 4, when the real fish only interacts with its most influential neighbor ($k_R = 1$). Neighbors are identified and ranked at each time step according to their instantaneous influence on the real fish, and the distributions are computed for neighbors of all influence ranks. Probability density functions (PDFs) of (**A**–**C**) the inter-individual distance $d$ and (**D**–**F**) the viewing angle $\psi$.

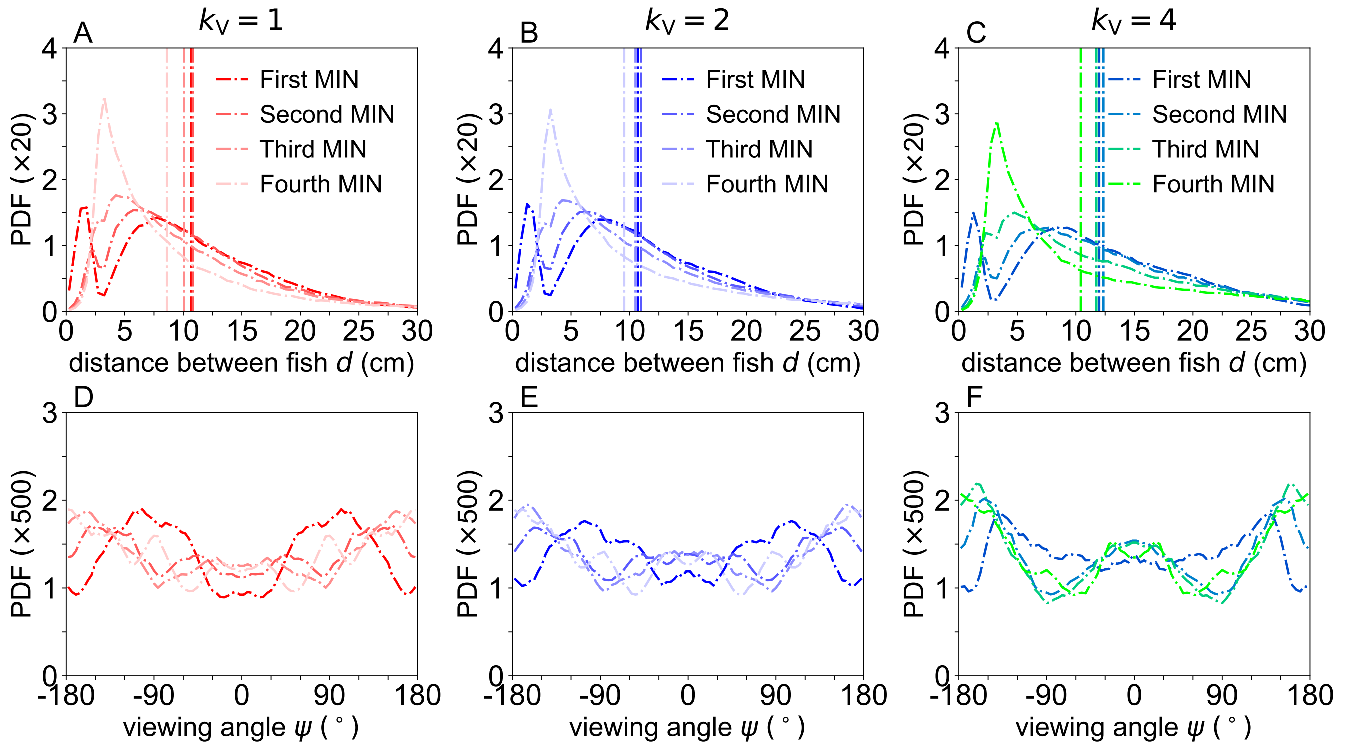


FIG. S9. **Spatial distribution of influential neighbors of the real fish ranked by influence.** Relative positions of the most influential neighbors (MIN) of the real fish measured in the experiments for the three social interaction strategies of the virtual fish, $k_V = 1$, 2, and 4, when the real fish interacts only with its two most influential neighbors ($k_R = 2$). Neighbors are identified and ranked at each time step according to their instantaneous influence on the real fish, and the distributions are computed for neighbors of all influence ranks. Probability density functions (PDFs) of (**A**–**C**) the inter-individual distance $d$ and (**D**–**F**) the viewing angle $\psi$.

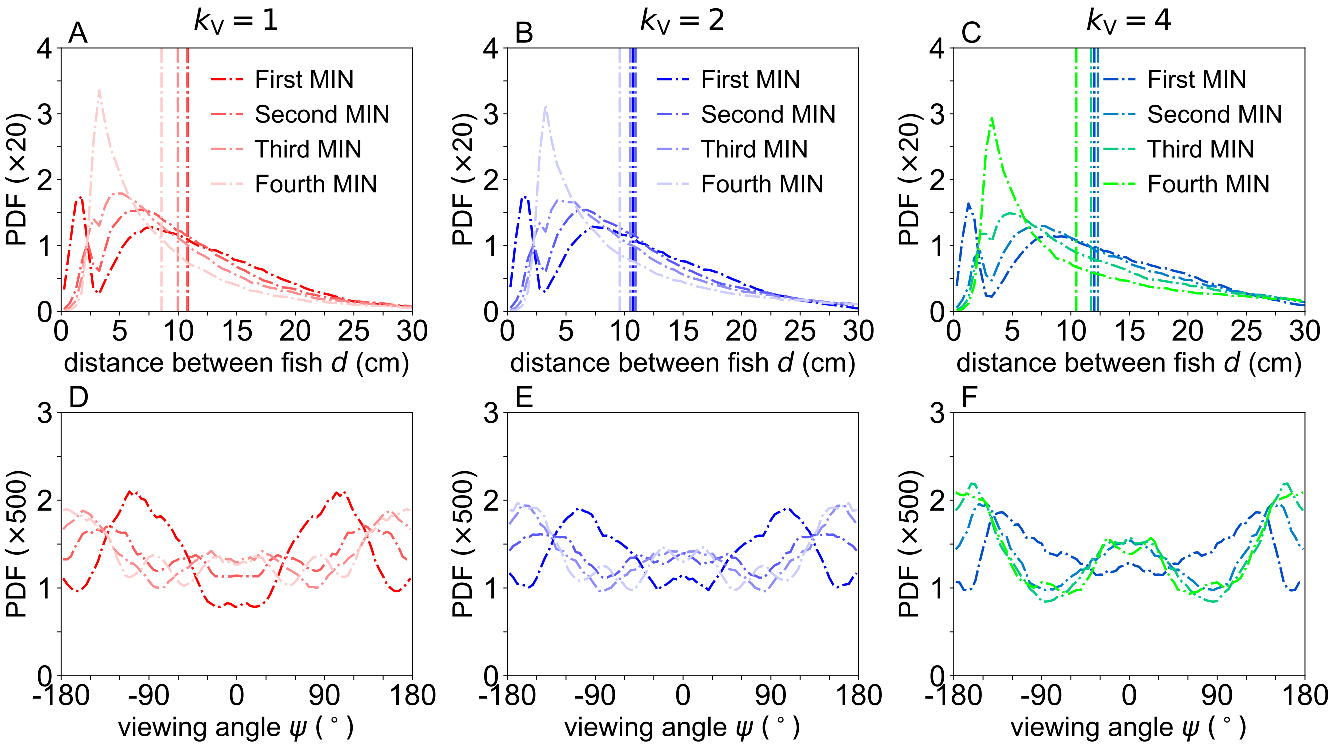


FIG. S10. **Spatial distribution of influential neighbors of the real fish ranked by influence.** Relative positions of the most influential neighbors (MIN) of the real fish measured in the experiments for the three social interaction strategies of the virtual fish, $k_V = 1$, 2, and 4, when the real fish when the real fish interacts with all its neighbors ($k_R = 4$). Neighbors are identified and ranked at each time step according to their instantaneous influence on the real fish, and the distributions are computed for neighbors of all influence ranks. Probability density functions (PDFs) of (**A**–**C**) the inter-individual distance $d$ and (**D**–**F**) the viewing angle $\psi$.

### SUPPLEMENTAL TABLES

TABLE S1. **Model parameters for different social interaction strategies.** Parameter values and multiplicative coefficients used in the simulations for each social interaction strategy, corresponding to different numbers of interacting neighbors.

| Multiplicative $k$-depending coefficient | Symbol | $k = 1$ | $k = 2$ | $k = 4$ |
|---|---|---|---|---|
| Preferred speed $v_0$ | $C_{\mathrm{v}}^{(0)}$ | 3 | 12 | 16 |
| Wall force coefficient | Coef_f_w | 0.5 | 1.9 | 4 |
| Attraction coefficient | Coef_f_att | 5 | 1.5 | 0.6 |
| Alignment coefficient | Coef_f_ali | 10 | 3 | 0.9 |

| Parameter or multiplicative coefficient | Symbol | Value for all $k$ |
|---|---|---|
| Parallel noise | $\sigma_{\parallel}$ | 4.5 |
| Perpendicular noise | $\sigma_{\perp}$ | 3.5 |
| Vertical noise $\sigma_z$ | $\sigma_z$ | 2.6 |
| Noise correlation time (s) | $\tau_{xy}$, $\tau_z$ | 0.15 |
| Attraction range | $C_{f_{\mathrm{Att}},d}^{(1)}$ | 40 |
| Preferred depth $z_0$ | $C_{f_z}^{(0)}$ | 5 |
| Effective distance between fish head and position coordinates (cm) | $d_c$ | 2.5 |
| Random angular correction after rejection | $d_{\mathrm{bounce}}$ | 0.25 |
| Threshold of short-range repulsion (cm) | d_c_Att | 4 |
| Intensity of short-range repulsion | f_c | 2 |
| Horizontal friction coefficient | Coef_Friction | 0.25 |
| Vertical friction coefficient | Coef_Friction_z | 1 |
| Speed adaptation | Coef_Adapt | 0 |
| Rotational force coefficient | Coef_f_rot | 1 |
| Vertical force coefficient | Coef_f_z | 1 |
| Vertical attraction coefficient | Coef_f_att_z | 1 |

TABLE S2. **Hellinger distance (×10).** Hellinger distance between experimental and numerical probability density functions for each behavioral metric and modeling strategy. All values have been scaled (×10) to facilitate comparison: values below 1 (in bold) indicate high statistical similarity.

| STRATEGY | $\mathbf{v}$ | $\boldsymbol{r}_\mathbf{w}$ | $\boldsymbol{\theta}_\mathbf{w}$ | $\boldsymbol{d}$ | $\boldsymbol{d}_\mathbf{NN}$ | $\boldsymbol{\psi}$ | $\boldsymbol{\Delta\phi}$ | $\boldsymbol{\Delta\phi}_\mathbf{NN}$ | $\langle$**All**$\rangle$ |
|---|---|---|---|---|---|---|---|---|---|
| $k_\mathrm{V} = 1$, $k_\mathrm{R} = 1$ | **0.80** | **0.95** | **0.70** | 1.00 | **0.72** | **0.16** | **0.77** | **0.63** | **0.72** |
| $k_\mathrm{V} = 1$, $k_\mathrm{R} = 2$ | **0.73** | 1.05 | **0.59** | **0.74** | **0.50** | **0.19** | **0.41** | **0.31** | **0.57** |
| $k_\mathrm{V} = 1$, $k_\mathrm{R} = 4$ | **0.74** | **0.89** | **0.59** | **0.76** | **0.61** | **0.22** | **0.13** | **0.26** | **0.52** |
| $k_\mathrm{V} = 2$, $k_\mathrm{R} = 1$ | **0.86** | 1.68 | **0.45** | **0.87** | **0.71** | **0.09** | **0.93** | **0.68** | **0.78** |
| $k_\mathrm{V} = 2$, $k_\mathrm{R} = 2$ | **0.78** | 1.93 | **0.89** | 1.22 | **0.92** | **0.22** | 1.49 | 1.19 | 1.08 |
| $k_\mathrm{V} = 2$, $k_\mathrm{R} = 4$ | **0.84** | 1.86 | 1.02 | **0.77** | **0.58** | **0.20** | 1.24 | 1.08 | **0.95** |
| $k_\mathrm{V} = 4$, $k_\mathrm{R} = 1$ | **0.48** | 1.38 | **0.48** | **0.93** | **0.73** | **0.35** | **0.75** | **0.56** | **0.71** |
| $k_\mathrm{V} = 4$, $k_\mathrm{R} = 2$ | **0.55** | 1.57 | **0.72** | 1.28 | **0.90** | **0.13** | 1.33 | 1.11 | **0.95** |
| $k_\mathrm{V} = 4$, $k_\mathrm{R} = 4$ | **0.67** | 1.66 | 1.00 | 1.38 | **0.97** | **0.27** | 1.48 | 1.36 | 1.10 |

TABLE S3. **Summary statistics of behavioral observables across experimental and simulation conditions.** Mean ± standard deviation of the different observables measured in experiments and in model simulations for all investigated conditions. For each condition, statistics were computed by averaging across all individuals.

| STRATEGY | $\boldsymbol{P}$ | $\boldsymbol{M}$ | $\mathbf{v}$(cm/s) | $\boldsymbol{r}_\mathbf{w}$(cm) | $\boldsymbol{d}$(cm) | $\boldsymbol{d}_\mathbf{NN}$(cm) | $\boldsymbol{v}_\mathbf{z}$(cm/s) | $\boldsymbol{z}$(cm) |
|---|---|---|---|---|---|---|---|---|
| $k_\mathrm{V} = 1,\ k_\mathrm{R} = 1$ | $0.74 \pm 0.28$ | $0.42 \pm 0.26$ | $12.14 \pm 4.45$ | $8.00 \pm 4.79$ | $8.11 \pm 5.06$ | $4.43 \pm 2.95$ | $0.00 \pm 0.82$ | $4.25 \pm 0.96$ |
| $k_\mathrm{V} = 1,\ k_\mathrm{R} = 2$ | $0.71 \pm 0.28$ | $0.45 \pm 0.27$ | $11.88 \pm 4.74$ | $8.06 \pm 4.84$ | $8.68 \pm 5.36$ | $4.75 \pm 3.17$ | $0.00 \pm 0.82$ | $4.25 \pm 0.96$ |
| $k_\mathrm{V} = 1,\ k_\mathrm{R} = 4$ | $0.67 \pm 0.28$ | $0.46 \pm 0.27$ | $11.69 \pm 4.86$ | $8.54 \pm 4.83$ | $9.31 \pm 5.57$ | $5.08 \pm 3.32$ | $0.00 \pm 0.82$ | $4.26 \pm 0.96$ |
| $k_\mathrm{V} = 2,\ k_\mathrm{R} = 1$ | $0.84 \pm 0.21$ | $0.47 \pm 0.26$ | $10.84 \pm 4.61$ | $7.02 \pm 4.65$ | $7.65 \pm 4.59$ | $4.18 \pm 2.73$ | $0.00 \pm 0.82$ | $4.21 \pm 0.87$ |
| $k_\mathrm{V} = 2,\ k_\mathrm{R} = 2$ | $0.88 \pm 0.17$ | $0.48 \pm 0.26$ | $10.73 \pm 4.50$ | $6.51 \pm 4.46$ | $7.14 \pm 4.21$ | $3.93 \pm 2.54$ | $0.00 \pm 0.82$ | $4.19 \pm 0.84$ |
| $k_\mathrm{V} = 2,\ k_\mathrm{R} = 4$ | $0.86 \pm 0.19$ | $0.51 \pm 0.26$ | $10.79 \pm 4.43$ | $6.61 \pm 4.47$ | $7.72 \pm 4.74$ | $4.23 \pm 2.88$ | $0.00 \pm 0.82$ | $4.19 \pm 0.83$ |
| $k_\mathrm{V} = 4,\ k_\mathrm{R} = 1$ | $0.86 \pm 0.18$ | $0.59 \pm 0.26$ | $11.13 \pm 4.25$ | $6.41 \pm 4.12$ | $8.34 \pm 5.15$ | $4.50 \pm 3.04$ | $0.00 \pm 0.83$ | $4.18 \pm 0.82$ |
| $k_\mathrm{V} = 4,\ k_\mathrm{R} = 2$ | $0.89 \pm 0.14$ | $0.62 \pm 0.25$ | $11.32 \pm 4.09$ | $5.78 \pm 3.85$ | $7.71 \pm 4.71$ | $4.17 \pm 2.82$ | $0.00 \pm 0.82$ | $4.15 \pm 0.77$ |
| $k_\mathrm{V} = 4,\ k_\mathrm{R} = 4$ | $0.90 \pm 0.13$ | $0.66 \pm 0.23$ | $11.45 \pm 3.95$ | $5.51 \pm 3.56$ | $7.52 \pm 4.63$ | $4.06 \pm 2.77$ | $0.00 \pm 0.82$ | $4.13 \pm 0.74$ |
| $k_\mathrm{V} = 1$ (exp) | $0.65 \pm 0.27$ | $0.44 \pm 0.27$ | $11.45 \pm 4.36$ | $8.42 \pm 4.64$ | $9.45 \pm 6.59$ | $4.98 \pm 3.80$ | $0.00 \pm 0.96$ | $4.36 \pm 1.07$ |
| $k_\mathrm{V} = 2$ (exp) | $0.73 \pm 0.23$ | $0.45 \pm 0.27$ | $9.99 \pm 4.39$ | $8.00 \pm 4.47$ | $8.65 \pm 5.89$ | $4.66 \pm 3.69$ | $0.00 \pm 1.01$ | $4.52 \pm 1.21$ |
| $k_\mathrm{V} = 4$ (exp) | $0.77 \pm 0.21$ | $0.57 \pm 0.27$ | $11.05 \pm 4.09$ | $6.46 \pm 3.78$ | $9.47 \pm 6.57$ | $4.92 \pm 4.13$ | $-0.01 \pm 0.99$ | $4.54 \pm 1.29$ |

TABLE S4. **Summary statistics of variables measured separately for real and virtual fish in experiments.** Mean ± standard deviation of the measured variables under the different experimental conditions. For each condition, statistics for the real fish are computed from a single individual, whereas statistics for the virtual fish are obtained by averaging across all virtual individuals.

| STRATEGY | $\mathbf{v}$(cm/s) | $\boldsymbol{r}_\mathbf{w}$(cm) | $\boldsymbol{\theta}_\mathbf{w}^+$(°) | $\boldsymbol{\theta}_\mathbf{w}^-$(°) | $\boldsymbol{d}$(cm) | $\boldsymbol{d}_\mathbf{NN}$(cm) | $\boldsymbol{z}$(cm) | $\boldsymbol{d}_\mathbf{z}$(cm) | $\boldsymbol{\Delta\phi}$(°) |
|---|---|---|---|---|---|---|---|---|---|
| $k_\mathrm{V} = 1$ (real) | $12.5 \pm 4.6$ | $7.5 \pm 4.4$ | $88.4 \pm 38.4$ | $-88.4 \pm 40.0$ | $10.0 \pm 6.7$ | $5.5 \pm 4.2$ | $4.51 \pm 1.30$ | $-0.19 \pm 1.46$ | $-0.9 \pm 84.2$ |
| $k_\mathrm{V} = 1$ (virtual) | $11.2 \pm 4.3$ | $8.7 \pm 4.7$ | $90.8 \pm 40.6$ | $-90.9 \pm 40.6$ | $9.3 \pm 6.6$ | $4.9 \pm 3.7$ | $4.32 \pm 1.00$ | $0.05 \pm 1.35$ | $0.2 \pm 81.2$ |
| $k_\mathrm{V} = 2$ (real) | $11.7 \pm 4.5$ | $7.6 \pm 4.4$ | $88.2 \pm 38.7$ | $-88.2 \pm 40.8$ | $10.4 \pm 7.0$ | $6.6 \pm 5.7$ | $4.93 \pm 1.69$ | $-0.51 \pm 1.62$ | $-0.5 \pm 87.4$ |
| $k_\mathrm{V} = 2$ (virtual) | $9.6 \pm 4.3$ | $8.1 \pm 4.5$ | $91.5 \pm 35.0$ | $-92.3 \pm 38.4$ | $8.2 \pm 5.5$ | $4.2 \pm 2.8$ | $4.42 \pm 1.02$ | $0.13 \pm 1.29$ | $0.1 \pm 67.8$ |
| $k_\mathrm{V} = 4$ (real) | $12.3 \pm 4.6$ | $5.9 \pm 3.7$ | $89.0 \pm 32.3$ | $-88.7 \pm 38.7$ | $11.6 \pm 7.8$ | $7.4 \pm 6.6$ | $4.99 \pm 1.99$ | $-0.57 \pm 1.79$ | $-3.6 \pm 80.9$ |
| $k_\mathrm{V} = 4$ (virtual) | $10.7 \pm 3.9$ | $6.6 \pm 3.8$ | $90.5 \pm 21.9$ | $-91.2 \pm 27.3$ | $8.9 \pm 6.1$ | $4.3 \pm 2.9$ | $4.42 \pm 1.01$ | $0.14 \pm 1.24$ | $0.9 \pm 62.1$ |

TABLE S5. **Hellinger distance (×10).** Hellinger distance between real fish and virtual fish probability density functions for each behavioral metric and modeling strategy in experiments. All values have been scaled (×10) to facilitate comparison: values below 1 (in bold) indicate high statistical similarity.

| STRATEGY | $\mathbf{v}$ | $\boldsymbol{r}_{\mathbf{w}}$ | $\boldsymbol{\theta}_{\mathbf{w}}$ | $\boldsymbol{d}$ | $\boldsymbol{\psi}$ | $\boldsymbol{\Delta\phi}$ | $\langle$**All**$\rangle$ |
|---|---|---|---|---|---|---|---|
| $k_{\mathrm{V}} = 1$ | 1.33 | 1.03 | **0.37** | **0.61** | **0.53** | **0.70** | **0.76** |
| $k_{\mathrm{V}} = 2$ | 1.81 | **0.65** | **0.65** | 1.32 | **0.55** | 1.67 | 1.11 |
| $k_{\mathrm{V}} = 4$ | 1.53 | 1.17 | 1.70 | 1.42 | **0.56** | 1.44 | 1.30 |

### SUPPLEMENTAL MOVIES

**Movie S1** Closed-loop bio-hybrid interactions between a real fish and four virtual conspecifics when each virtual fish only interacts with its most influential neighbor ($k_V = 1$). Video excerpt of an experiment in which a real fish interacts with the anamorphic projections of four virtual conspecifics whose motion is controlled in real time by the model, enabling closed-loop interactions both among virtual fish and with the real fish. In this condition, each virtual fish only interacts with its most influential neighbor ($k_V = 1$). Top left: user interface allowing real-time visualization of the trajectories of the real fish (in red) and the four virtual fish (in the $xy$ and $xz$ planes) and on-the-fly modification of the parameters of the model driving the virtual fish. Bottom left: real-time 3D tracking of the real fish. Right panel: anamorphic rendering of the four virtual fish projected onto the bowl by the rendering application according to the 3D position of the real fish.

**Movie S2** Closed-loop bio-hybrid interactions between a real fish and four virtual conspecifics when each virtual fish interacts with its two most influential neighbors ($k_V = 2$). Video excerpt of an experiment in which a real fish interacts with the anamorphic projections of four virtual conspecifics whose motion is controlled in real time by the model, enabling closed-loop interactions both among virtual fish and with the real fish. In this condition, each virtual fish interacts only with its two most influential neighbors ($k_V = 2$). Top left: user interface allowing real-time visualization of the trajectories of the of the real fish (in red) and the four virtual fish (in the $xy$ and $xz$ planes) and on-the-fly modification of the parameters of the model driving the virtual fish. Bottom left: real-time 3D tracking of the real fish. Right panel: anamorphic rendering of the four virtual fish projected onto the bowl by the rendering application according to the 3D position of the real fish.

**Movie S3** Closed-loop bio-hybrid interactions between a real fish and four virtual conspecifics when each virtual fish interacts with all four neighbors ($k_V = 4$). Video excerpt of an experiment in which a real fish interacts with the anamorphic projections of four virtual conspecifics whose motion is controlled in real time by the model, enabling closed-loop interactions both among virtual fish and with the real fish. In this condition, each virtual fish interacts only with its two most influential neighbors ($k_V = 2$). Top left: user interface allowing real-time visualization of the trajectories of the real fish (in red) and the four virtual fish (in the $xy$ and $xz$ planes) and on-the-fly modification of the parameters of the model driving the virtual fish. Bottom left: real-time 3D tracking of the real fish. Right panel: anamorphic rendering of the four virtual fish projected onto the bowl by the rendering application according to the 3D position of the real fish.

**Movie S4** Experimental bio-hybrid fish group dynamics with increasing social coupling of the virtual-fish. Representative sequences extracted from experiments showing the collective dynamics of a bio-hybrid fish group composed of one real fish (red) interacting with four virtual conspecifics (blue) across three successive conditions differing in the number of influential neighbors considered by the virtual fish. The video presents, in sequence, minimal social coupling when each virtual fish responds only to its single most influential neighbor ($k_V = 1$), intermediate coupling when virtual fish respond to their two most influential neighbors ($k_V = 2$), and strong coupling when each virtual fish responds to all group members ($k_V = 4$). The sequences illustrate real-time closed-loop interactions and how progressive changes in social information filtering shape emergent collective motion.

**Movie S5** Simulated bio-hybrid fish group dynamics with increasing social integration of the real-fish while virtual fish only interact with their single most influential neighbor ($k_V = 1$). Numerical simulations of a bio-hybrid fish group composed of one real fish (red) and four virtual conspecifics (blue) shown across three successive conditions differing in the number of influential neighbors integrated by the simulated real fish while virtual fish only interact with their single most influential neighbor ($k_V = 1$). The video presents, in sequence, the case where both real and virtual fish respond to a single influential neighbor ($k_R = 1$), followed by conditions where the simulated real fish responds to two neighbors ($k_R = 2$), and to all four neighbors ($k_R = 4$).

**Movie S6** Simulated bio-hybrid fish group dynamics with increasing social integration of the real-fish while virtual fish interact with their two most influential neighbors ($k_V = 2$). Numerical simulations of a bio-hybrid fish group composed of one real fish (red) and four virtual conspecifics (blue) shown across three successive conditions in which virtual fish interact with their two most influential neighbors ($k_V = 2$) while the number of neighbors integrated by the simulated real fish varies. The video presents, in sequence, the case where the simulated real fish responds to a single influential neighbor ($k_R = 1$), followed by conditions where the simulated real fish responds to two neighbors ($k_R = 2$), and to all four neighbors ($k_R = 4$).

**Movie S7** Simulated bio-hybrid fish group dynamics with increasing social integration of the real-fish while virtual fish interact with all group members ($k_V = 4$). Numerical simulations of a bio-hybrid fish group composed of one real fish (red) and four virtual conspecifics (blue) shown across three successive conditions in which virtual fish respond to all group members as influential neighbors ($k_{\mathrm{V}} = 4$) while the number of neighbors integrated by the simulated real fish varies. The video presents, in sequence, the case where the simulated real fish responds to a single influential neighbor ($k_{\mathrm{R}} = 1$), followed by conditions where it responds to two neighbors ($k_{\mathrm{R}} = 2$), and where both real and virtual fish respond to all group members ($k_{\mathrm{R}} = 4$).
